# Profiling of the human intestinal microbiome and bile acids under physiologic conditions using an ingestible sampling device

**DOI:** 10.1101/2022.01.19.476920

**Authors:** Dari Shalon, Rebecca Neal Culver, Jessica A. Grembi, Jacob Folz, Peter Treit, Les Dethlefsen, Xiandong Meng, Eitan Yaffe, Sean Spencer, Handuo Shi, Andrés Aranda-Díaz, Andrew D. Patterson, George Triadafilopoulos, Susan P. Holmes, Matthias Mann, Oliver Fiehn, David A. Relman, Kerwyn Casey Huang

## Abstract

The spatiotemporal structure of the human microbiome and metabolome reflects and determines regional intestinal physiology and may have implications for disease. Yet, we know little about the distribution of microbes and their products in the gut because of reliance on stool samples and limited access only to some regions of the gut using endoscopy in fasting or sedated individuals. To address these deficiencies, we developed and evaluated a safe, ingestible device that collects samples from multiple regions of the human intestinal tract during normal digestion. The collection of 240 intestinal samples from 15 healthy individuals using the device revealed significant differences between microbes and metabolites present in the intestines versus stool. Certain microbial taxa were differentially enriched, and bile acid profiles varied along the intestines and were highly distinct from those of stool. Correlations between gradients in bile acid concentrations and microbial abundance predicted species that altered the bile acid pool through deconjugation. Overall, we identified heterogeneous intestinal profiles of bacterial taxa and metabolites indicating that non-invasive multi-regional sampling of the intestinal tract under physiological conditions can help elucidate the roles of the gut microbiome and metabolome in human physiology and disease.

## Introduction

The human intestinal tract harbors the vast majority of the microbes residing in or on our bodies^1^; their genetic content and biochemical transformation capabilities are hundreds of times larger than those encoded by the human genome^2^. Humans depend on our gut microbes for food digestion, immune system regulation, and protection against pathogens, among other critical functions^3^. An important yet often overlooked aspect of the gut is its regional heterogeneity and differentiation. Due to the difficulties in accessing and sampling the human intestinal tract, stool has been the main source of information for human gut microbiome studies^4^. However, stool reflects the waste products and downstream effluent of the gut, within which regional variation is lost. For example, key metabolites such as bile acids are altered upstream by microbial transformations and then partially absorbed by the host before excretion in stool^5^. The regions of the gut distal to the stomach (duodenum, jejunum, ileum, and colon) differ dramatically in nutrient availability, pH, oxygen partial pressure, mucosal structure, and flow rates^6^. As a result, distinct microbial communities with specialized functions and immune niches are thought to be present in each intestinal region^7^. Indeed, the microbial communities at mucosal sites in the large intestine are known to be distinct from those of stool^8^. Moreover, the host proteome in mice segregates more strongly by intestinal location than by colonization state^9^, underscoring the potential for regional differences in host gene expression. Thus, a true understanding of how gut microbes impact human physiology and vice versa requires local sampling of the gut microbiome and its chemical environment in an unperturbed state.

Historically, sampling from the human intestinal tract without disturbance or contamination has been challenging and results have at times been contradictory^10^. In a recent study, we discovered substantial regional variability in microbiota composition across spatial scales of only a few inches throughout the intestines of deceased organ donors^11^. However, organ donors have been typically treated with antibiotics prior to organ harvesting, and even in cases in which the intestinal tract has been sampled immediately following cessation of life support, it is typically ischemic or necrotic^11^. Duodenal sampling from live subjects using upper endoscopy has a high probability of inadvertent contamination from oral, esophageal, or gastric contents. Endoscopic access to the mid-jejunum requires a 2-3 h procedure involving general anesthesia or sedation and the procedure is performed in fasting subjects^12,13^. Alternatively, a stoma created by exteriorization of the ileum through the abdominal wall can provide a ready source of intestinal samples, but this procedure is invasive and reflects altered gut anatomy and physiology^14^. While the intestinal tracts of model organisms such as mice can be invasively sampled, the pH profiles, peristalsis, diet, physiology, gastrointestinal disorders, and key metabolites such as bile acids^15^ differ markedly between humans and animals^16^, making human studies more informative and most relevant to human physiology and disease.

Bile acids are major chemical components of the human intestinal tract and are critical for food digestion, lipid absorption, host signaling and neurohormonal regulation of diverse physiological processes. Bile acids have been implicated in a wide range of disorders including inflammatory bowel disease^17^, metabolic disorders^17^, and neurological diseases^18,19^. Primary bile acids are synthesized from cholesterol in the liver^5^ where they are conjugated to glycine or taurine to form bile salts^20^. Bile salts are secreted into the duodenum and transformed extensively by bacteria in the intestinal tract into secondary bile acids through deconjugation, dehydroxylation, and epimerization reactions^21^. Approximately 95% of bile acids are actively transported through the distal ileal epithelium into the portal vein back to the liver^5^ where they are transformed back into bile salts and re-secreted into the duodenum multiple times during the course of a single meal. Biles acids interact with both bacteria and the host; bile acids are toxic to many bacteria at high concentrations due to membrane disruption^22^, and act as host signaling molecules by activating a variety of receptors in several human cell types^23,24^. Different receptors generally bind distinct subsets of bile acid chemical structures^21,23^. Despite their important effects on the microbiome and their signaling properties, we know little about the chemical diversity and concentrations of bile acids within the intestines; instead, human studies to date have relied on the measurement of non-representative subsets of bile acids found in stool or blood.

To measure intestinal bile acid and microbial profiles during normal digestion, we developed and evaluated a capsule device that samples the luminal contents of the small intestine and ascending colon. We observed distinct microbial communities present in the intestines, as compared to stool. Using mass spectrometry-based metabolomics, we discovered gradients of microbially transformed bile acids along the intestinal tract. We also identified correlations between the concentrations of microbially modified bile acids and the abundance of specific gut bacterial species. These discoveries reveal novel biology that is not accessible from the study of stool or endoscopic sampling of the intestinal tract.

## Results

### Development of a noninvasive capsule device for sampling the human intestines

The sampling capsule is a single-use, passive device that collects fluid from the human intestines for *ex vivo* analysis. The device contains a collapsed collection bladder capped by a one-way valve inside a dissolvable capsule with an enteric coating (Fig. 1A). The enteric coating prevents contact between the collection bladder and the surrounding environment prior to entry into the intestines. The pH of the intestines typically rises from 4-6 in the duodenum to 7-8 in the ileum^25^. Once the device reaches a pre-set pH level sufficient to dissolve the enteric coating, the collection bladder expands and draws in luminal contents through the one-way valve. To sample from four distinct regions of the intestinal tract, four devices were ingested as a set after a subject ate a meal of their choosing, wherein each device type in a set was designed to open at different, progressively higher pH levels. Device type 4 included a time-delay coating to bias collection toward the ascending colon where the pH typically drops relative to the terminal ileum^25^ (Methods, Fig. 1A). Each device collects up to 400 µL of luminal contents, rather than the mucosal or epithelial-associated habitat, motivated by the observations that bacterial density is higher in the lumen than within the mucosa^26^, that most of the mucosal-associated bacteria are represented in the luminal contents^27^, and that many metabolites of interest are in the lumen. After the bladder fills, the one-way valve prevents further entrance of liquid. The ingested devices are recovered from the stool and the collected samples are extracted for analysis. Thus, our device provides unique potential for multi-region collection of microbes and metabolites within the intestines during normal digestion.

**Figure 1:**
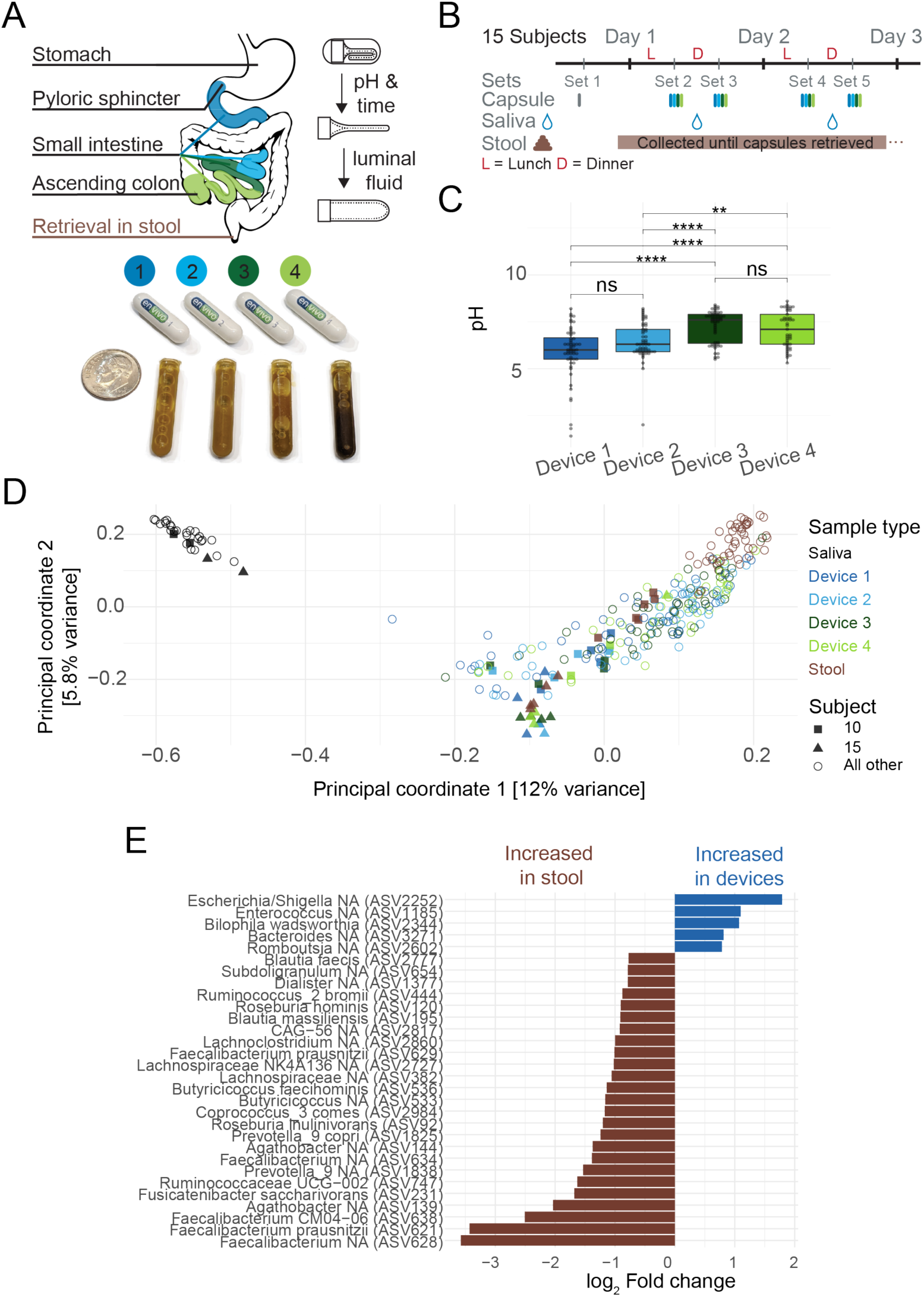
Capsule devices enable longitudinal sampling of the human intestine. A) Overview of the intended sampling locations (left) of the four types of sampling devices (bottom right) in packaged form for ingestion and as full collection bladders containing intestinal samples as collected from the stool. (Top right) Device contains a folded bladder capped with a one-way valve within an enterically coated capsule; the enteric coating dissolves once the designated pH has been reached, enabling the bladder to unfold and draw in up to 400 µL of luminal fluid. A U.S. dime is included for scale. B) Study design and timeline for the collection of samples from 15 healthy adult subjects. C) The pH of the contents in devices designed to open at locations spanning the proximal to distal intestinal tract. There is a trend along the intestinal tract. Points represent individual capsules. ns: not significant, *: *p*≤0.05, **: *p*≤0.01, ****: *p*≤0.0001, Bonferroni-adjusted Wilcoxon rank sum test. D) Principal coordinates analysis based on Canberra distance between microbial communities using 16S rRNA gene amplicon sequence variants (ASVs, *n*=455) with ≥3 reads in ≥5% of samples. The analysis highlights separation of communities by sample type. Read counts were log_2_-transformed. Each point represents an individual sample and is colored by the sample type (stool, saliva, and device types 1-4). Filled squares and triangles identify two outlier subjects (#10 and #15) who had taken oral antibiotics in the 5 months prior to intestinal sampling. E) ASVs with a minimum log_2_-fold change between devices and stool of 0.75 were detected. Only ASVs that were significantly differentially abundant (*p*<0.05 after Benjamini-Hochberg correction) are shown.

We first sought to confirm whether the devices could be targeted to specific intestinal locations in humans and would progress through the intestinal tract without contamination. In a feasibility study, we connected devices targeting the jejunum and ascending colon to a capsule endoscope and visualized successful *in vivo* sampling in a human subject (Movie S1). To assess the potential effects of incubation of the samples while the device transited the gut, we retrieved a set of 4 devices from a single bowel movement 32 h after ingestion and immediately incubated them in an anaerobic chamber at 37 °C to simulate the effect of extended retention in the intestinal tract. Aliquots from each sample at the start of incubation (32 h), and at hours 58 and 87 were subjected to 16S rRNA gene amplicon sequencing. The rank abundance of the 30 most abundant amplicon sequence variants (ASVs, a proxy for species) at 32 h shifted after 58 h by a median of 8-16 ranks and after 87 h by 12-30 ranks (Fig. S1). The 9-17 ASVs that increased from below to above the detection limit during incubation accounted for a relative abundance of 9.4%-13.8% after 58 h and 5.2%-18% after 87 h, presumably due to growth during incubation. Thus, while outgrowth can potentially alter our assessments of microbiota composition, major changes are not expected for transit times of ∼58 h or less. Within these experimental limitations, we demonstrate below that microbes and metabolites display a gradient along the intestine and are highly distinct from stool samples.

### Spatially distinct microbial communities detected along the longitudinal axis of the human intestinal tract

To assess compositional and functional differences within the intestinal microbiome, we carried out a clinical study with 15 healthy human subjects. First, a single device was swallowed and retrieved to ensure that no complications arose during passage of the device through the gut (set 1, Fig. 1B); the contents of these devices were not analyzed. Subsequently, sets of 4 devices (each device type within a set having a different enteric coating) were ingested twice daily (3 h after lunch and 3 h after dinner) on two consecutive days, for a total of 4 sets (sets 2-5, Fig. 1B). Each set of 4 devices was designed to generate a profile along the intestinal tract. All subjects consumed their normal diets and kept a food log. All devices safely exited all subjects and were successfully retrieved. No adverse events were reported. Contemporaneous stool and saliva samples were also collected regularly (Fig. 1B).

Of the 240 devices, 218 collected >50 µL of intestinal fluids and were subjected to 16S rRNA gene and metagenomic sequencing; the remainder sampled <50 µL or filled with gas, most likely from the colon. Of the 218 devices that sampled >50 µL, we obtained sufficient sequencing reads from 210 samples (Fig. S2, Methods), which were thus the focus of subsequent analyses, along with sequencing data from saliva and stool samples. The pH profiles of the samples collected by the four devices types (Fig. 1C) reasonably matched previously published measurements of pH values in the human intestines, with a general increase in pH from the proximal to distal region of the intestines followed by a pH drop in the ascending colon^25^. The time between device ingestion and recovery ranged from 8 to 67 h (Fig. S3A), consistent with previous reports of broadly distributed transit times^28^. Given typical gastric emptying times and the 3 h post-meal interval before devices were swallowed, the devices likely entered the small intestines with the final contents of the preceding meal^29–31^. Nonetheless, the contents of the subsequent meal were more strongly associated with gut transit time of the devices (Fig. S3B,C).

A principal coordinates analysis identified variation of microbiota composition along the intestinal tract and across disparate anatomical regions (saliva, intestines, and stool). Saliva samples were significantly segregated from intestinal and stool samples across all subjects (PERMANOVA *p*=0.001, Fig. 1D), indicating that the contents of all devices did not reflect the composition of the oral microbiota. Furthermore, we identified 2 subjects (#10 and #15, Fig. 1D) whose stool, and to some degree intestinal, samples clustered separately. Upon follow-up questioning, these subjects reported taking antibiotics within the past 1 (#10) and 5 months (#15). When considering each subject individually, 23%±10% (137±70 of 582±85) of the ASVs detected in the devices were not detected in the subject’s saliva or stool; the median relative abundance of these 137 ASVs was low (<0.4%). Similarly, 12%±8% of the ASVs in stool were not detected in the subject’s intestinal samples and the median relative abundance of these ASVs was low (<0.6%) in all but one outlier subject (#3) whose intestinal samples were dominated by a single species. In line with previous studies^32^, we observed higher relative abundances of the Proteobacteria phylum in the intestinal tract compared to stool (Fig. S4), including a *Bilophila wadsworthia* ASV, consistent with previous reports of *B. wadsworthia*’s key role in the small intestine^33,34^. Four additional ASVs, from the *Escherichia*/*Shigella, Enterococcus, Bacteroides*, and *Romboutsia* genera, were also significantly more abundant (adjusted *p*<0.05 and log_2_-fold change >0.75) in intestinal samples than stool (Fig. 1E). The *Romboutsia* genus was named in 2014 following isolation of a species from rat ileal digesta^35^, consistent with this genus having a niche in the small intestine. In contrast, 30 ASVs were more abundant in stool than intestinal samples (Fig. 1E).

We observed more intra-subject microbial variability among intestinal samples than among stool or saliva samples (Fig. 2A), suggesting that the devices collect from a more heterogenous habitat. While each device type was designed to sample from a specific region of the intestinal tract, comparisons of microbiota composition among devices of the same type but swallowed at different times are potentially masked by variability in meal contents, periprandial neurohormonal variations, intestinal motility, pH, or the intestinal microbiota itself. We assessed technical and biological variability by having one subject ingest 4 devices of the same type simultaneously; this procedure was repeated twice for each of the device types 1-4 over the course of 2 months. Devices of the same type ingested at the same time revealed more similar microbial communities than devices of the same type ingested at different times (Fig. 2B). The increased variance in microbiota composition due to this temporal variability is comparable to the variance due to spatial variability along the intestine, as assessed using sets of 4 devices of distinct types ingested at the same time (Fig. 2B). Moreover, intestinal samples (unlike saliva or stool) were often dominated by a single ASV with relative abundance >40% (Fig. 2D). Consequently, individual intestinal samples contained communities with lower alpha diversity relative to the intra-subject diversity represented by all samples from a device of a certain type, or from sets of all samples from devices swallowed at the same time (Fig. 2C,D, Fig. S5). Thus, much of the higher variability across intestinal samples relative to stool is likely due to the dynamic and heterogeneous nature of the microbiota along the intestinal tract.

**Figure 2:**
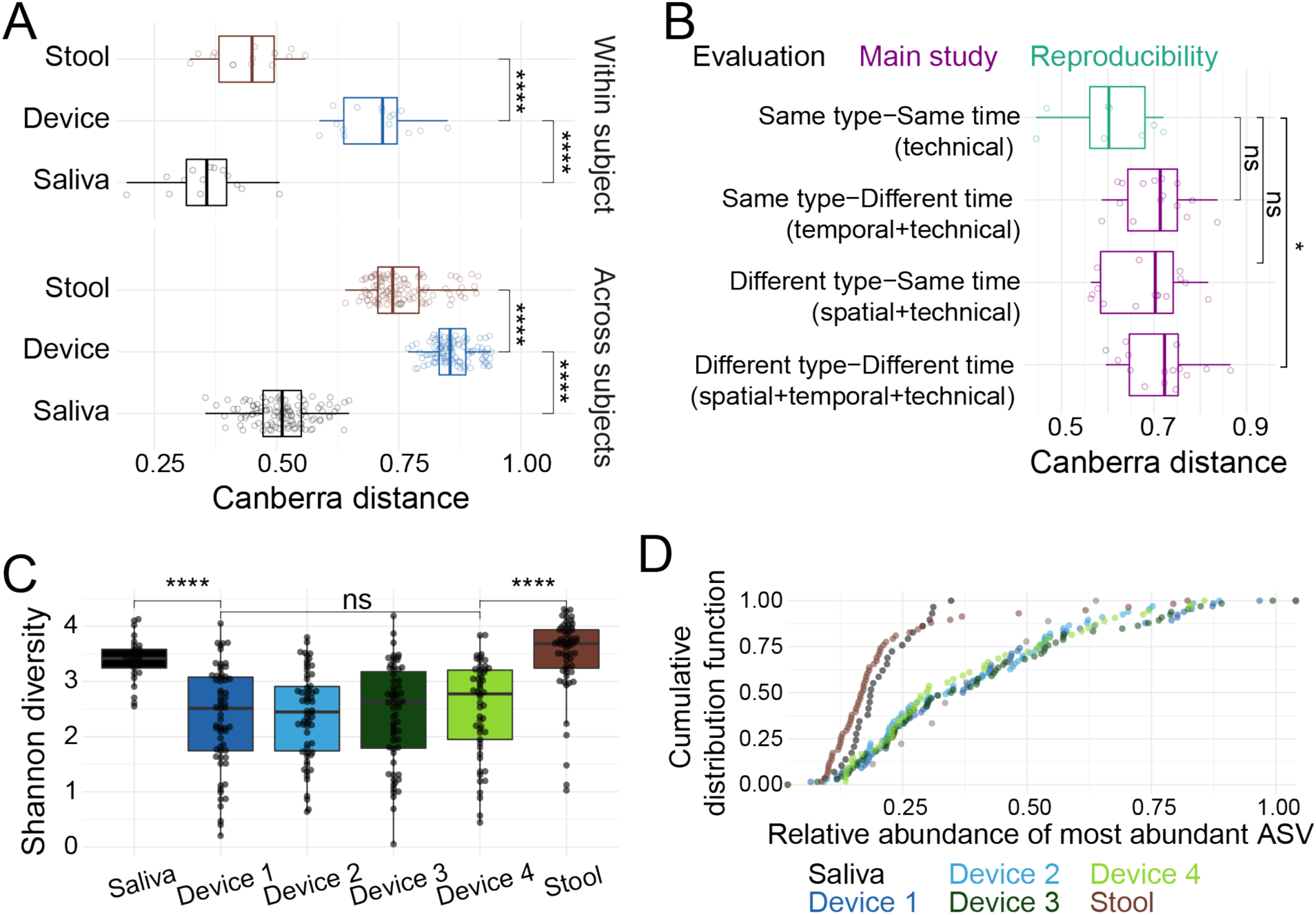
Microbiota variation across capsule types suggests patchy structure. A) Microbiota composition varied significantly more between intestinal samples that it did between stool or between saliva samples. This variation persisted both within (top) and across (bottom) subjects. Each point in the top panel is the mean pairwise Canberra distance between all samples for a subject. In the bottom panel, each point is the mean of all pairwise comparisons between all samples from any two subjects. B) Temporal, spatial, and technical variability in microbiota composition of intestinal samples (purple) were higher than in technical replicates (green), in which one subject swallowed 4 of the same device type simultaneously (the subject did so twice for each of the 4 device types). Each point represents the mean pairwise Canberra distance between intestinal samples from the same subject. Canberra distances for (A,B) were computed from log_2_-transformed read counts of 16S rRNA gene amplicon sequence variants (ASVs, *n*=455) with read count ≥3 in ≥5% of samples. Microbial communities from devices of the same type ingested at the same time were more similar than devices of the same type ingested at different times, although this was not statistically robust to Bonferroni correction (Wilcoxon rank sum adjusted *p*=0.058) given the small number of observations. C) The Shannon diversity of saliva and stool samples was higher than that of intestinal samples. Each point is a single sample. D) Capsules were more likely to be dominated by a single ASV as compared with stool or saliva. Each point is a single sample. ns: not significant, *: *p*≤0.05, ****: *p*≤0.0001, Bonferroni-corrected Wilcoxon rank sum test.

### Distinct patterns of bile acids along the human intestinal tract

Glycine- and taurine-conjugated forms of the primary bile acids cholic acid (CA) and chenodeoxycholic acid (CDCA) are secreted from the liver and gallbladder into the duodenum and are then subjected to various microbial transformations (Fig. 3A) that could be expected to lead to longitudinal bile acid gradients along the intestinal tract. To quantify bile acid profiles, we performed targeted LC/MS-MS metabolomics with multiple reaction monitoring (MRM) on seventeen bile acids from the supernatants of all intestinal samples and from stool. Both the total concentration of bile acids and their relative levels in the intestinal samples were highly variable (Fig. 3B), yet distinct trends were observed. The total concentration of bile acids generally decreased ∼2 fold in samples collected by type 4 devices and ∼10 fold in stool relative to samples collected by type 1 devices (Fig. 3B), likely reflecting active reabsorption of bile acids along the intestines^5^.

**Figure 3:**
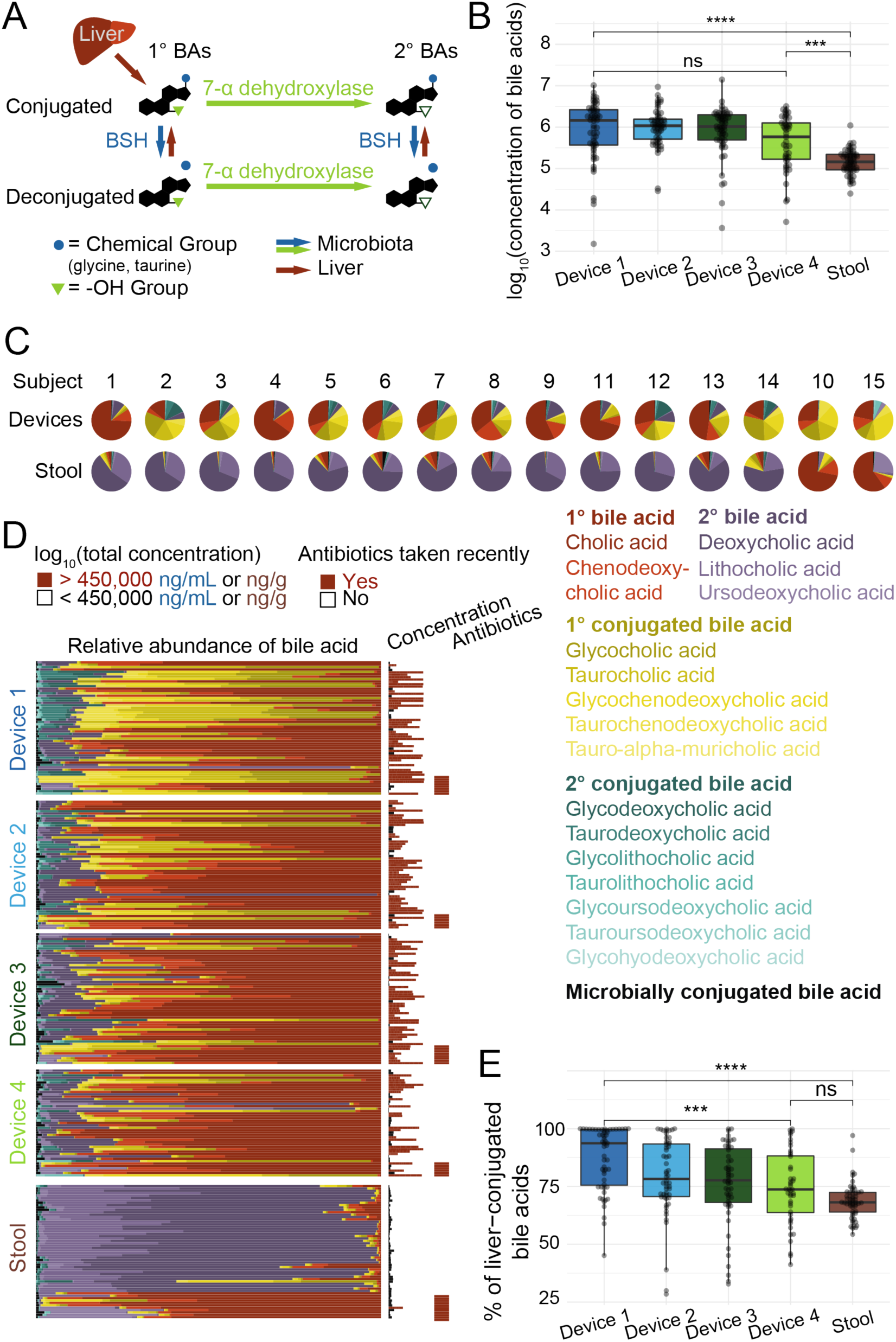
Capsules capture different bile acid profiles along the intestinal tract as compared to stool. A) Schematic of bile acid modifications by the liver and microbiota. The liver releases bile acids conjugated with glycine or taurine. Dehydroxylation by gut microbes converts primary (1°) to secondary (2°) bile acids. Microbial bile salt hydrolases (BSHs) deconjugate amino acids from bile salts. B) Total concentration of all bile acids decreased along the intestinal tract. Shown are log_10_-transformed concentrations in units of ng/mL or ng/g for intestinal or stool samples, respectively. Boxplots show the median, 25th, and 75th quartiles. ns: not significant, *: *p*≤0.05, ****: *p*≤0.0001, Bonferroni-corrected Wilcoxon rank sum test. C) All subjects showed distinct bile acid profiles in intestinal compared with stool samples. Deoxycholic and lithocholic acid dominated the stool, but not the intestines, in all but two subjects (#10 and #15). Profiles are means of bile acid relative concentrations over all samples in the respective categories (intestinal or stool) per subject. D) Bile acid profiles in each sample, grouped by sample type, demonstrate variability throughout the intestinal tract. Profiles are the relative abundance of bile acid chemical structures. Histogram on the right displays the total concentration (ng/mL or ng/g for intestinal samples and stool, respectively) of all bile acids in each sample, and bars (far right) indicate whether the subject had taken antibiotics in the past 5 months. E) Percent of liver-conjugated bile acids decreased significantly along the intestinal tract. Boxplots show the median, 25th, and 75th quartiles. *: *p*≤0.05, ****: *p*≤0.0001, Bonferroni-corrected Wilcoxon rank sum test.

The stool bile acid profiles of two subjects (#10 and #15) were similar to their intestinal samples with a dominant fraction of CA (Fig. 3C, S8), in contrast to all other subjects. These are the two subjects that reported recently taking antibiotics and exhibited substantially different microbiota composition from the other subjects (Fig. 1D). All intestinal and stool samples from subjects 10 and 15 also lacked DCA and LCA (Fig. 3C), suggesting that the microbes necessary for the dehydroxylation reaction required to produce these bile acids may have been eliminated by the use of antibiotics.

In all other subjects, the relative levels and various bile acid classes differed dramatically between the intestinal and stool samples. Intestinal samples were mostly dominated by the primary bile acid CA while stool samples were dominated by the secondary bile acid deoxycholic acid (DCA) (Fig. 3C,D), likely due to the prolonged exposure of bile acids to microbial enzymes in the colon. These trends in bile-acid profiles across device types provide further support that the sampling locations of the devices were longitudinally distributed along the intestinal tract, and indicate that stool-based measurements of bile acids do not reflect the true composition of bile acids along the intestinal tract.

### Detection of gradients of bile acid modifications along the intestinal tract

Bile acids are modified in the intestinal tract by microbial enzymes that deconjugate glycine or taurine or remove hydroxyl group(s) from the sterol backbone (Fig. 3A). Deconjugation is performed by bile salt hydrolases (BSHs), which cleave glycine and taurine from the bile acid backbone, and BSH homologs are present in ∼25% of bacterial strains found previously in metagenomic sequencing of human stool stamples^36^. Although neither the abundance nor the genetic diversity of BSH genes was distinct among device types or between intestinal and stool samples based on metagenomic sequencing in this study (Fig. S6), we observed a significant monotonic decrease in the percentage of glycine- and taurine-conjugated bile acids from device type 1 to device type 4 samples (Fig. 3E). These results quantify a trend of increasing deconjugation along the intestinal tract and into stool (Fig. 3E)^37^.

Dehydroxylation reactions require several enzymes to transform primary to secondary bile acids and are thought to be predominantly active in the low redox state of the colon^37^. Consistent with the majority of dehydroxylation occurring in the large intestine, we found that secondary bile acids did not change substantially across device types, but significantly increased in stool samples, which were dominated by secondary unconjugated bile acids (Fig. 3D, S7, S8). The presence of secondary bile acids found in intestinal samples are likely due to dehydroxylation of primary bile acids in the small intestines or re-introduction of secondary bile acids present in bile into the duodenum; the presence of secondary bile acids in bile is expected given previous evidence of their presence at ∼25% of bile acids secreted from the gallbladder^37^. In sum, the variation we detected in bile acid profiles throughout the intestinal tract demonstrates the regionality of the microbial activity and biochemical environment of the intestines, further highlighting the limitations of relying on stool for microbiome and bile acid studies.

The significantly different bile acid profiles in intestinal compared to stool samples indicate that it is unlikely that stool contaminated the intestinal sampling devices during transit or sample recovery. However, due to the large increase in microbial density along the intestinal tract^37^, stool contamination could affect microbiota composition. We therefore used a statistical approach to identify samples as potentially contaminated based on microbial community composition (Methods). Re-analysis of all data after removing a subset of samples that displayed any signal of potential contamination from stool (*n*=38) resulted in the same statistical trends as the complete group of samples.

### Linking specific microbes to bile acid deconjugation and dehydroxylation

We next sought to exploit the variation in conjugated bile acid concentrations across intestinal samples to identify candidate bacterial species responsible for deconjugation. Given the decrease in the percent of liver-conjugated bile acids across capsule types (Fig. 3E), we reasoned that the abundance of microbial taxa most responsible for deconjugation might be inversely correlated with the concentration of conjugated bile acids, even against the background of potential regulation of deconjugation by the host or antimicrobial activity of bile acids.

We focused on primary bile acids, which dominate the pool of conjugated bile acids, namely glycocholic acid (GCA) and taurocholic acid (TCA). The concentration of both GCA and TCA decreased across device types and was significantly lower in stool (Fig. 4A,B). GCA concentration was negatively correlated with the relative abundance of *Anaerostipes hadrus* and *Faecalibacterium prausnitzii* (Fig. 4C). Similarly, the relative abundance of *Bilophila wadsworthia* and *Alistipes putredinis* exhibited a statistically significant negative correlation with TCA concentration (Fig. 4D). Across all subjects, we obtained 440 high-quality metagenome assembled genomes (MAGs, completeness >90% and contamination <10%, dereplicated to 99% ANI) and searched for the canonical BSH gene in each using a hidden Markov model. We found putative BSH genes in *Anaerostipes hadrus* (7 of 8 MAGs) and *Alistipes putredinis* (4 of 4 MAGs), in accordance with previous literature^38^. By contrast, none of the 12 *Faecalibacterium prausnitzii* MAGs nor the 3 *Bilophila wadsworthia* MAGs contained any putative BSH genes, suggesting these taxa may utilize glycine and taurine^33^ generated by other microbial deconjugation reactions.

**Figure 4:**
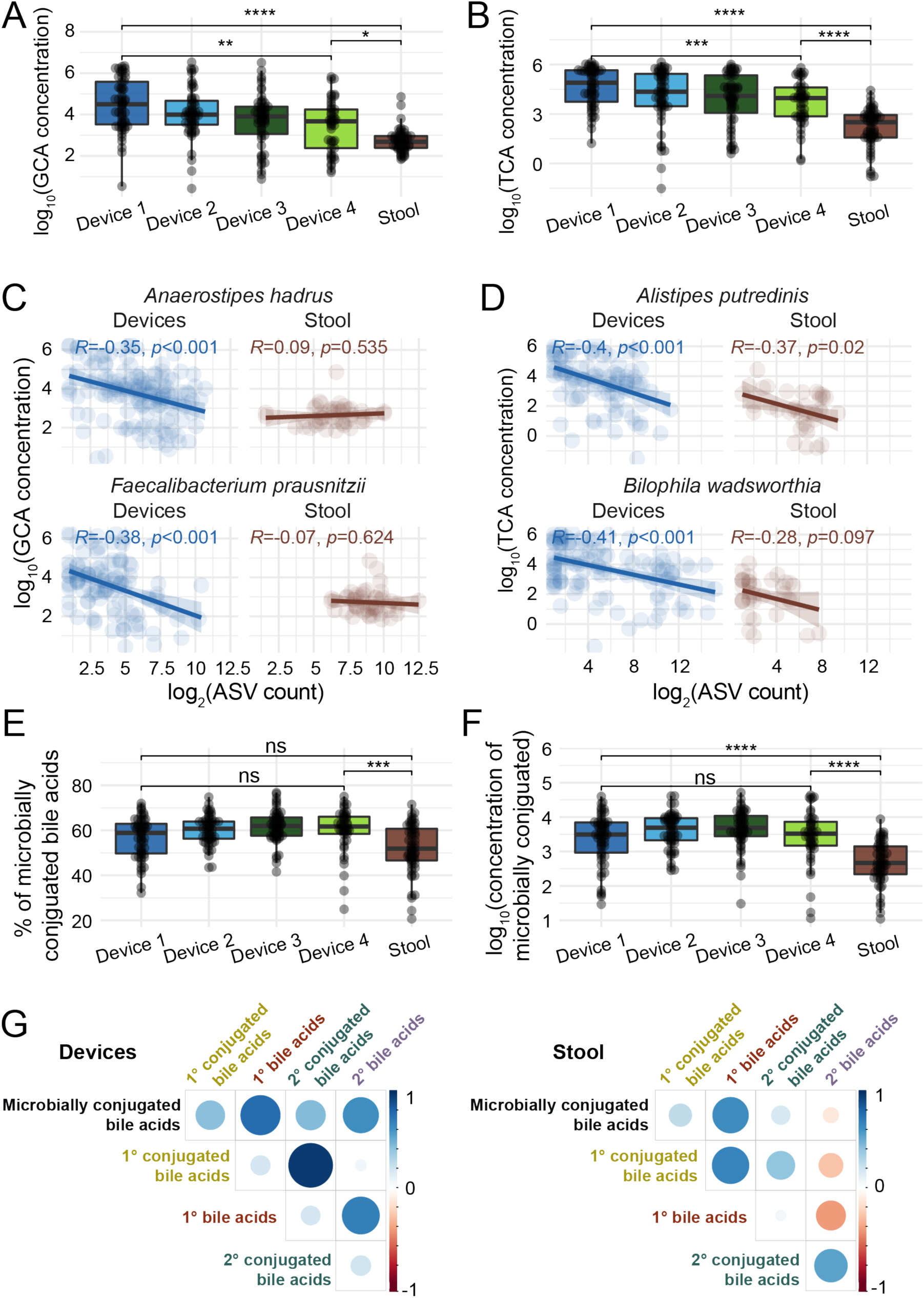
Microbial bile acid modifications detected in the intestinal tract. A,B) Glycocholic acid (GCA, (A)) and taurocholic acid (TCA, (B)) concentration decreased along the intestinal tract. Shown are log_10_-transformed concentrations in units of ng/mL or ng/g for intestinal or stool samples, respectively. C,D) The log_2_(ASV count) of an *Anaerostipes hadrus* and a *Faecalibacterium prausnitzii* ASV was significantly negatively correlated (Pearson) with the concentration of GCA (C), and the log_2_(ASV count) of an *Alistipes putredinis* and a *Bilophila wadsworthia* ASV was significantly negatively correlated with TCA concentration (D). Only ASVs with *p*<0.01 after a Benjamini-Hochberg correction are shown. These correlations were much weaker or not significant in the stool samples. E) The percent of microbially conjugated bile acids increased along the intestinal tract and was significantly higher in intestinal samples (particularly type 4 devices) compared with stool. F) The concentration of microbially conjugated bile acids was significantly higher in intestinal compared to stool samples. The concentration did not differ significantly across device types. Approximate concentrations in ng/mL or ng/g for intestinal or stool samples, respectively. G) Correlations between bile acid classes differed between the intestinal and stool samples. Shown are Pearson’s correlation coefficients using log_10_-transformed concentrations in ng/mL or ng/g for intestinal or stool samples, respectively. All boxplots show the median, 25^th^, and 75^th^ quartiles. Points are individual intestinal or stool samples. ns: not significant, **: *p*≤0.01, ****: *p*≤0.0001, Bonferroni-corrected Wilcoxon rank sum test. Read counts were log_2_-transformed.

Concentrations of taurochenodeoxycholic acid (TCDCA) were also negatively correlated with *B. wadsworthia* and *A. putredinis* log_2_(abundance), and TDCA was negatively correlated with *B. wadsworthia* abundance, indicating that these species likely interact with various taurine-conjugated bile acids (Fig. S9). We focused mainly on *B. wadsworthia* since it was differentially abundant in intestinal samples compared with stool (Fig. 1E). The name of the *Bilophila* genus reflects its growth stimulation by high concentrations of bile^39^, and work in mice based on stool and gall bladder extracts has linked high-fat diets with high levels of TCA and *B. wadsworthia*, potentially due to the ability of *B. wadsworthia* to use taurine for growth^33^. Importantly, in stool, the relative abundance of these ASVs was correlated only weakly or not at all with TCA concentration (Fig. 4D), indicating that the devices enable identification of correlations between bile acids and microbes that would not be evident from stool.

### Microbially conjugated bile acid concentrations vary along the intestinal tract

Recently, bile acids conjugated to amino acids other than glycine and taurine (e.g., tyrosocholic acid (TyroCA), leucocholic acid (LeuCA), phenylalanocholic acid (PhenylCA) were discovered in the gut of mice and humans^20^. Synthesis of these newly discovered bile acid conjugates has been reported to be carried out by microbes in the intestinal tract, and their levels have been hypothesized to differ significantly between healthy and diseased states^20^. Using untargeted LC-MS/MS analysis with data-dependent MS/MS acquisition, we detected these novel bile acids, along with 18 additional amino acid-bile acid conjugates in various hydroxylation forms across 13 amino acids in the intestinal samples of all subjects (Table S3). Microbially conjugated bile acids were at significantly higher concentration and accounted for a significantly higher fraction of the bile acid pool in intestinal samples compared with stool, and the fraction increased from type 1 devices to type 4 devices (Fig. 4E,F).

We also found that the concentrations of primary and secondary liver-conjugated bile acids were correlated with each other, while the concentration of microbially conjugated bile acids was strongly correlated with that of deconjugated bile acids across intestinal samples (Fig. 4G, S7). These findings emphasize the effect of different anatomical regions and routes of formation and degradation for liver-conjugated bile acids (glycine and taurine conjugates) and microbially conjugated bile acids. In stool, the concentration of microbially conjugated bile acids was correlated with the concentration of primary deconjugated bile acids and inversely correlated with the concentration of secondary deconjugated bile acids (Fig. 4E), highlighting major differences in the metabolite environment of the intestines versus stool.

## Discussion

To date, studies of the human gut microbiota and bile acids have relied mainly on stool. In this study, enabled by the development and implementation of an ingestible sampling device, we demonstrated that analysis of stool does not provide a complete or accurate representation of the longitudinal and temporal variability of the microbiota composition and bile acid contents within the intestines. Microbiota and bile acid profiles were distinct along the intestines and from other disparate regions, including saliva and stool. We also discovered correlations between bile acid profiles and microbial taxa in the intestines of healthy humans, suggesting distinct functional capabilities of the intestinal microbiota compared to stool, and quantified the distribution of recently discovered microbially conjugated bile acids throughout the gut. Moreover, in subjects who had recent antibiotic exposure, bile acid profiles of stool suggested dramatic alterations compared to intestinal samples. Regional and repetitive sampling of gradients along the intestine during digestion is thus required to capture the variable nature of the gut microbiome and the production, transformation, and utilization of key metabolites.

One limitation of our study is that the exact location of sample collection within the intestine could not be clearly defined or validated. Variability in intestinal peristalsis and pH during normal digestion may cause capsule devices within a set to experience different pH trajectories, hence they may open before or after their intended collection sites. Despite this limitation, analysis of 210 intestinal samples from 15 individuals discovered consistent trends of biochemical and microbial activity in the human intestines. More consistent sampling along a longitudinal gradient might be attained in future studies by collecting multiple sequential samples into a single capsule device in a timed manner.

The wide variability among intestinal samples across our cohort of 15 subjects and within each subject at different time points highlights the need for future studies utilizing larger cohorts and longer-term sampling. We envision interrogating how diet and disease differentially influence the intestinal microbiota and metabolome. Indeed, measurements from the proximal intestinal microbial ecosystem will be critical for future clinical studies of spatially restricted human intestinal diseases and therapeutic interventions directed at these disorders.

Nonetheless, our study demonstrates the feasibility and potential of a safe and non-invasive method for collection, characterization, and quantification of the intestinal microbiota and bile acids along the human intestinal tract during normal digestion. This new capability, when deployed at scale, should improve our understanding of the dynamic and intertwined nature of human metabolic pathways with our resident gut microbes, and their potential involvement in normal physiology and disease.

## Methods

### Ingestible capsule sampling device

The capsule sampling device (CapScan^®^, Envivo^®^ Bio Inc, San Carlos CA) consists of a one-way valve capping a hollow elastic collection bladder. The device is prepared for packaging by evacuating the collection bladder, folding it in half, and packaging the folded device inside a dissolvable capsule measuring 6.5 mm in diameter and 23 mm long, onto which an enteric coating is applied. The capsule and the enteric coating prevent contamination of the collection bladder from oral-pharyngeal and gastric microbes during ingestion. When the device reaches the target pH, the enteric coating and capsule disintegrate. The target pH is pH 5.5 for type 1, pH 6 for type 2, pH 7.5 for type 3 and type 4, with type 4 also having a time delay coating to bias collection toward the ascending colon. After the enteric coating disintegrates, the collection bladder unfolds and expands into a tube 6 mm in diameter and 33 mm long, thereby drawing in up to 400 µL of gut luminal contents through the one-way valve. The one-way valve maintains the integrity of the sample collected inside the collection bladder as the device moves through the colon and is exposed to stool.

In this study, subjects concurrently ingested sets of 4 capsules, each with distinct coatings to target the proximal to medial regions of the small intestine (coating types 1 and 2) and more distal regions (coating types 3 and 4). After sampling, the devices were passed in the stool into specimen-collection containers and immediately frozen. After completion of sampling, the stool was thawed and the devices were retrieved by study staff. The elastic collection bladders were rinsed in 70% isopropyl alcohol and punctured with a sterile hypodermic needle attached to a 1-mL syringe for sample removal. Samples were transferred into microcentrifuge tubes and the pH was measured with an InLab Ultra Micro ISM pH probe (Mettler Toledo). A 40-µL aliquot was spun down for 3 min at 10,000 rcf, and its supernatant was used for metabolomics analysis. The rest of the sample was frozen until being thawed for DNA extraction.

### Study design

The study was approved by the WIRB-Copernicus Group IRB (study #1186513) and informed consent was obtained from each subject. Healthy volunteers were selected to exclude participants suffering from clinically detectable gastrointestinal conditions or diseases that would potentially interfere with data acquisition and interpretation.

Subjects met all of the following criteria for study inclusion: (1) individuals between the ages of 18 and 70; (2) American Society of Anesthesiologists (ASA) physical status class risk of 1 or 2; (3) for women of childbearing potential, a negative urine pregnancy test within 7 days of screening visit, and willingness to use contraception during the entire study period; and (4) fluency in English, understands the study protocol, and is able to supply informed written consent, along with complying with study requirements.

Subjects with any of the following conditions or characteristics were excluded from the study: (1) history of any of the following: prior gastric or esophageal surgery, including lap banding or bariatric surgery, bowel obstruction, gastric outlet obstruction, diverticulitis, inflammatory bowel disease, ileostomy or colostomy, gastric or esophageal cancer, achalasia, esophageal diverticulum, active dysphagia or odynophagia, or active medication use for any gastrointestinal conditions; (2) pregnancy or planned pregnancy within 30 days from screening visit, or breast-feeding; (3) any form of active substance abuse or dependence (including drug or alcohol abuse), any unstable medical or psychiatric disorder, or any chronic condition that might, in the opinion of the investigator, interfere with conduct of the study; or (4) a clinical condition that, in the judgment of the investigator, could potentially pose a health risk to the subject while involved in the study.

Fifteen healthy subjects were enrolled in this study, and each swallowed at least 17 devices over the course of three days (for demographics, see Table S1). All ingested devices were recovered, and no adverse events were reported during the study. Of the 255 ingested devices, 15 were set 1 safety devices (not included in analysis) and 22 contained gas or low sample volume. An additional 8 of the 218 samples remaining did not provide sufficient sequencing reads (>2500 reads) to be included in further analysis (Fig. S2). Saliva samples were collected after evening meals and immediately frozen at -20 °C. Every bowel movement during the study was immediately frozen by the subject at -20 °C. Subject 1 provided additional samples for assessment of replicability and blooming. A total of 297 saliva, intestinal, and stool samples were analyzed.

### Blooming analysis

To assess the effect of in-body incubation on the contents of the devices between the time of sample collection and sample retrieval, a set of 4 devices (one of each type) was ingested by subject 1. Upon exit in a bowel movement at 32 h, the devices were immediately transferred to an anaerobic chamber and incubated at 37 °C. An aliquot of each sample was taken at 32 h (immediately after the bowel movement), 58 h, and 87 h (the latter two time points simulating lengthier gut transit times).

### DNA extraction and 16S rRNA gene sequence analysis

DNA was extracted using a Microbial DNA extraction kit (Qiagen)^40^ and 50 µL from a capsule device, 200 µL of saliva, or 100 mg of stool.

16S rRNA amplicons were generated using Earth Microbiome Project-recommended 515F/806R primer pairs and 5PRIME HotMasterMix (Quantabio 2200410) with the following program in a thermocycler: 94 °C for 3 min, 35 cycles of [94 °C for 45 s, 50 °C for 60 s, and 72 °C for 90 s], followed by 72 °C for 10 min. PCR products were cleaned, quantified, and pooled using the UltraClean 96 PCR Cleanup kit (Qiagen 12596-4) and Quant-iT dsDNA High Sensitivity Assay kit (Invitrogen Q33120). Samples were sequenced with 250-bp reads on a MiSeq instrument (Illumina).

Sequence data were de-multiplexed using the Illumina bcl2fastq algorithm at the Chan Zuckerberg BioHub Sequencing facility. Subsequent processing was performed using the R statistical computing environment (v. 4.0.3)^41^ and DADA2 as previously described using pseudo-pooling^42^. truncLenF and truncLenR parameters were set to 250 and 180, respectively. Taxonomy was assigned using the Silva rRNA database v132^43^. Samples with >2500 reads were retained for analyses. A phylogenetic tree was constructed using phangorn as previously described^44^. Shannon diversity was calculated using the phyloseq::estimate_richness function, which is a wrapper for the vegan::diversity function^45,46^. Since intestinal samples were often dominated by a single ASV (Fig. 2D), the Canberra distance metric was used for pairwise beta diversity comparisons. Only the 455 ASVs represented by ≥3 reads in ≥5% of samples were used to calculate distances.

### Contamination analysis

One concern in our study was potential downstream contamination of device samples with stool, even a small amount of which could alter the microbial community in the devices given the orders of magnitude higher concentration of bacteria in stool compared with the proximal intestines^37^. However, given the directional motility of the intestinal tract, one would expect intrinsic overlap between intestinal and stool microbial communities. Latent Dirichlet allocation with the topic models R package^47^ was used to identify co-occurring groups of microbes (’topics’^48^) from intestinal and stool samples for each subject. For each intestinal sample, the cumulative probability of topics identified as derived from the same subject’s stool was computed. Capsules with ≥10% of the total community identified as potentially originating from stool topics were flagged as possibly contaminated. Using this very conservative definition, 38 of the 210 intestinal samples with adequate sequencing read counts were identified as possibly contaminated. All analyses presented in this study used all available data to avoid bias, but all results were robust to the removal of samples identified as possibly contaminated.

### Metagenomic sequencing

Extracted DNA from the samples was arrayed in a 384-well plate and concentrations were normalized after quantification using the PicoGreen dsDNA Quantitation kit (ThermoFisher). DNA was added to a tagmentation reaction, incubated for 10 min at 55 °C, and immediately neutralized. Mixtures were added to 10 cycles of a PCR that appended Illumina primers and identification barcodes to allow for pooling of samples during sequencing. One microliter of each well was pooled, and the pooled library was purified twice using Ampure XP beads to select the appropriately sized bands. Finally, library concentration was quantified using a Qubit (Thermo Fisher). Sequencing was performed on a NovaSeq S4 instrument with read lengths of 2×146 bp.

### Pre-processing of raw sequencing reads and metagenomic assembly

Skewer v. 0.2.2^49^ was used to remove Illumina adapters, after which human reads were removed with Bowtie2 v. 2.4.1^50^. Metagenomic reads from a single saliva, intestinal, or stool sample were assembled with MEGAHIT v. 1.2.9^51^. Assembled contigs were binned with MetaBAT 2 v. 2.15^52^ into 7,655 genome bins. checkM v. 1.1.3^53^ and quast v. 5.0.2^54^ were used to assess quality; bins with >75% completeness and <25% contamination were dereplicated at 99% ANI (strain level) with dRep v. 3.0.0^55^, resulting in 696 representative metagenome assembled genomes (MAGs) across all samples. GTDB-Tk was used to assign taxonomy^56^. Default parameters were used for all computational tools.

### Sample preparation for LC-MS/MS analysis and bile acid quantification

Supernatants from intestinal samples were extracted using a modified 96-well plate biphasic extraction^57^. Samples in microcentrifuge tubes were thawed on ice and 10 µL were transferred to wells of a 2-mL polypropylene 96-well plate in a predetermined randomized order. A quality control (QC) sample consisting of a pool of many intestinal samples from pilot studies was used to assess analytical variation. QC sample matrix (10 µL) and blanks (10 µL of LC-MS grade water) were included for every 10th sample. One hundred seventy microliters of methanol containing UltimateSPLASH Avanti Polar Lipids (Alabaster, Alabama) as an internal standard were added to each well. Then, 490 µL of methyl-tert-butyl-ether (MTBE) containing internal standard cholesterol ester 22:1 were added to each well. Plates were sealed, vortexed vigorously for 30 s, and shaken on an orbital shaking plate for 5 min at 4 °C. The plate was unsealed and 150 µL of cold water were added to each well. Plates were re-sealed, vortexed vigorously for 30 s, and centrifuged for 12 min at 4000 rcf and 4 °C.

From the top phase of the extraction wells, two aliquots of 180 µL each were transferred to new 96-well plates, and two aliquots of 70 µL each from the bottom phase were transferred to two other new 96-well plates. Plates were spun in a rotary vacuum until dry, sealed, and stored at -80 °C until LC-MS/MS analysis. One of the 96-well plates containing the aqueous phase of extract was dissolved in 35 µL of HILIC-run solvent (8:2 acetonitrile/ water, v/v). Five microliters were analyzed using non-targeted HILIC LC-MS/MS analysis. Immediately after HILIC analysis, the 96-well plates were spun in a rotary vacuum until dry, sealed, and stored at -80 °C until targeted bile acid analysis.

Multiple dilutions were prepared for bile acid analysis as follows. The dried samples described above were dissolved in 60 µL of bile acid-run solvent (1:1 acetonitrile/ methanol (v/v) containing 6 isotopically labeled bile acid standards at 100 ng/mL) via 30 s of vortexing and 5 min of shaking on an orbital shaker. From this plate, 5 µL were transferred to a new 96-well plate and combined with 145 µL of bile acid-run solvent. Both dilutions were analyzed for all samples, and samples that still presented bile acids above the highest concentration of the standard curve (1500 ng/mL) were diluted 5:145 once more and re-analyzed. A 9-point standard curve that ranged from 0.2 ng/mL to 1500 ng/mL was used with all samples. The standard curve solutions were created by drying bile acid standard solutions to achieve the desired mass of bile acid standards and then dissolved in bile acid-run solvent. Three standard-curve concentration measurements were analyzed after every 20 samples during data acquisition along with one method blank.

Approximately 4 mg (±1 mg) of wet stool were transferred to 2-mL microcentrifuge tubes. Twenty microliters of QC mix were added to microcentrifuge tubes for QC samples. Blank samples were generated using 20 µL of LC-MS grade water. To each tube, 225 µL of ice-cold methanol containing internal standards (as above) were added, followed by 750 µL of ice-cold MTBE with CE 22:1. Two 3-mm stainless-steel grinding beads were added to each tube and tubes were processed in a Geno/Grinder automated tissue homogenizer and cell lyser at 1500 rpm for 1 min. One hundred eighty-eight microliters of cold water were then added to each tube. Tubes were vortexed vigorously and centrifuged at 14,000 rcf for 2 min. Two aliquots of 180 µL each of the MTBE layer and two aliquots of 50 µL each of the lower layer were transferred to four 96-well plates, spun in a rotary vacuum until dry, sealed, and stored at -80 °C until analysis with the intestinal samples. Stool samples were analyzed using HILIC non-targeted LC-MS/MS and diluted in an identical manner to intestinal samples as described above. Stool samples were analyzed in a randomized order after intestinal samples.

### Metabolomics data acquisition

Samples were analyzed using a Thermofisher Vanquish UHPLC system coupled to a Thermofisher TSQ Altis triple-quadrupole mass spectrometer. An Aquity BEH C18 column (1.7 µm, 2.1 mm×100 mm) with guard column Acquity BEH C18 (1.7 µm, 2.1 mm×5 mm) was used for chromatographic separation with mobile phases of A: LC-MS-grade water with 0.1% formic acid, and B: LC-MS-grade acetonitrile with 0.1% formic acid with a flow rate of 400 µL/min and column temperature of 50 °C. The gradient began at 20% B for 1 min, then shifted to 45% B between 1 and 11 min, then to 95% B between 11 and 14 min, then to 99% B between 14 and 14.5 min, 99% B was maintained until 15.5 min, then transitioned from 99% B to 20% B between 15.5 and 16.5 min, and maintained at 20% B until 18 min. Injection volume was 5 µL and MRM scans were collected for all bile acids and internal standards (Table S2).

### Metabolomics data processing

MRM scans were imported to Skyline^58^ software. Skyline performed peak integration for all analytes with given mass transitions and retention time windows (Table S2). The chromatogram for each analyte was manually checked to confirm correct peak integration. Peak area was exported for all analytes. Bile acid chemical structures were removed if there was not a convincing chromatographic peak observed in ≥1 sample. The ratio of analyte to its closest eluting internal standard was calculated and used for quantification. A linear model was fitted to standard curve points for each bile acid (*R*^2^>0.995 for all bile acids) and the model was applied to all samples and blanks to calculate concentrations. The average concentration reported for method blanks was subtracted from sample concentrations. Since multiple dilutions were analyzed for each sample, the measurement closest to the center of the standard curve (750 ng/mL) was used. Zero values were imputed with a concentration value between 0.001 and 0.1 ng/mL. Concentrations were reported as ng/mL for intestinal sample liquid supernatant, and ng/g for wet stool.

### Non-targeted bile acid quantification

Bile acids conjugated to amino acids (e.g., TyroCA, LeuCA, and PhenylCA) were not included in the list for targeted analysis. Nonetheless, 22 microbe-conjugated bile acids were detected during non-targeted data acquisition for intestinal and stool samples using HILIC chromatography as described previously^59^. Peaks corresponding to these microbially conjugated bile acids were annotated using *m*/*z* values for precursor mass, diagnostic MS/MS fragment ions (337.2526 for tri-hydroxylated and 339.2682 for di-hydroxylated bile acids), and the corresponding amide conjugate fragment ion, as reported previously^60^ (Table S3). MS/MS spectra from synthetic standards for three microbially conjugated bile acids (Fig. S10) served as positive controls based on previously collected experimental MS/MS spectra^20^. Non-targeted HILIC analysis did not include bile acid standard curves to allow for direct quantification, so approximate quantification was achieved by comparing the concentration of GCA from targeted analysis to GCA peak height intensity from non-targeted analysis. A quadratic model was fit to GCA values from both analyses (*R*^2^=0.89) and applied to the peak height intensity values of microbe-conjugated bile acids to calculate their approximate concentrations. Approximate concentrations were used for analysis of bile acids measured with non-targeted analysis.

## Supplementary Tables

**Table S1:**
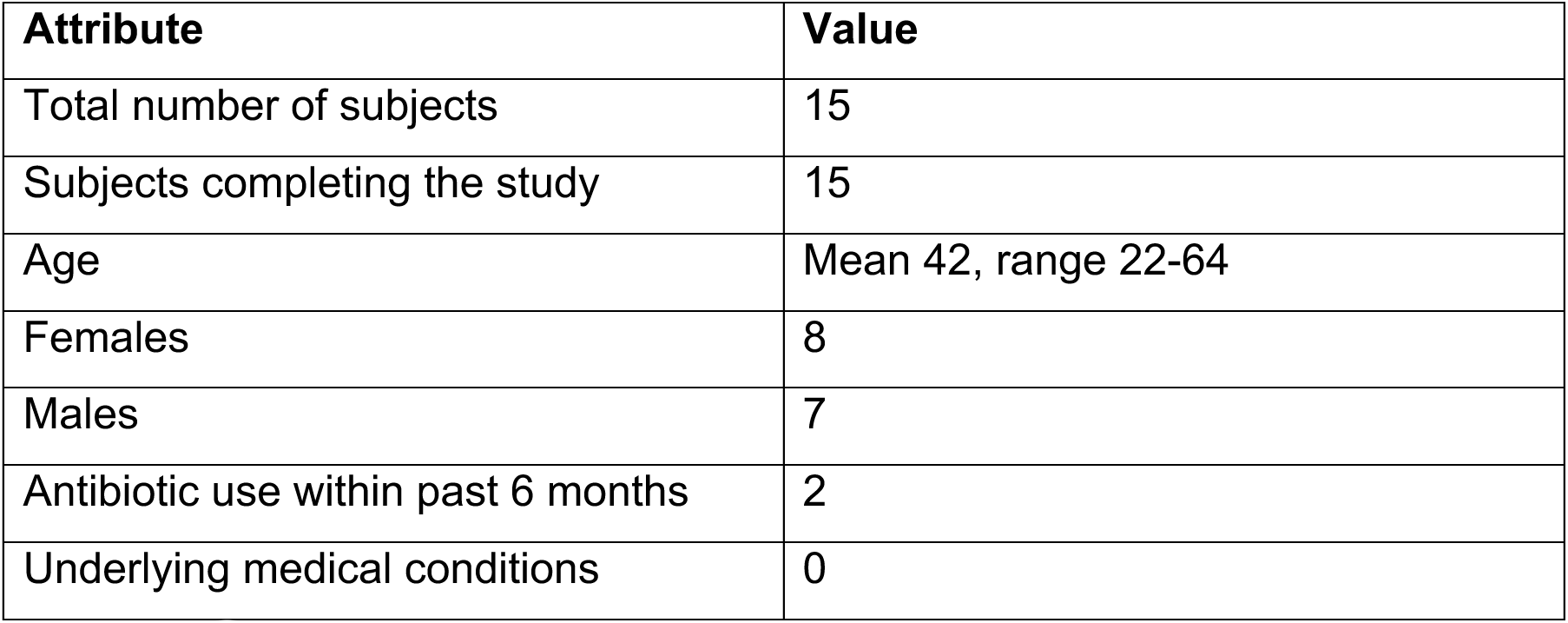
Subject demographics.

| Attribute | Value |
| --- | --- |
| Total number of subjects | 15 |
| Subjects completing the study | 15 |
| Age | Mean 42, range 22-64 |
| Females | 8 |
| Males | 7 |
| Antibiotic use within past 6 months | 2 |
| Underlying medical conditions | 0 |

**Table S2:** Bile acid quantification parameters. Parameters for LC-MS/MS data acquisition and data processing.

| Analyte | Precursor (m/z) | Product (m/z) | Collision energy (V) | Minimum dwell time (ms) | RF lens (V) | Source fragmentation | MRM start time (min) | MRM end time (min) | Analyte retention time (min) | RT integration window (± value) (min) |
| --- | --- | --- | --- | --- | --- | --- | --- | --- | --- | --- |
| 1-phenyl 3-hexadecanoic acid urea (PHAU) (internal standard) | 248.95 | 130.1 | 14.18 | 197.628 | 55 | 0 | 1.25 | 3.75 | 2.57 | 0.15 |
| Tauro-ω-muricholic acid | 514.475 | 80.042 | 55 | 72.825 | 152 | 0 | 2.5 | 7 | 3.35 | 0.08 |
| Tauro-α-muricholic acid | 514.475 | 80.042 | 55 | 72.825 | 152 | 0 | 2.5 | 7 | 3.53 | 0.08 |
| Tauro-β-muricholic acid | 514.475 | 80.042 | 55 | 72.825 | 152 | 0 | 2.5 | 7 | 3.72 | 0.15 |
| Tauroursodeoxycholic acid | 498.475 | 80.042 | 55 | 57.72 | 138 | 0 | 3.75 | 9.5 | 5.3 | 0.15 |
| Taurocholic acid | 514.475 | 80.042 | 55 | 72.825 | 152 | 0 | 2.5 | 7 | 5.68 | 0.15 |
| ω-Muricholic acid | 407.25 | 407.25 | 0 | 57.662 | 95 | 77.6 | 5.5 | 10.5 | 6.92 | 0.15 |
| Glycoursodeoxycholic acid | 448.288 | 74.054 | 33.18 | 57.72 | 93 | 69.4 | 5.5 | 8.5 | 7.03 | 0.08 |
| Glycohyodeoxycholic acid | 448.288 | 74.054 | 33.18 | 57.72 | 93 | 69.4 | 5.5 | 8.5 | 7.21 | 0.08 |
| Glycocholic acid | 464.425 | 74.054 | 35.92 | 57.72 | 143 | 0 | 6 | 8.5 | 7.22 | 0.15 |
| Glycocholic acid-d4 (internal standard) | 468.288 | 74.143 | 36.17 | 57.72 | 121 | 100 | 6 | 8.5 | 7.21 | 0.15 |
| α-muricholic acid | 407.25 | 407.25 | 0 | 57.662 | 95 | 77.6 | 5.5 | 10.5 | 7.24 | 0.1 |
| β-muricholic acid | 407.25 | 407.25 | 0 | 57.662 | 95 | 77.6 | 5.5 | 10.5 | 7.54 | 0.1 |
| Taurochenodeoxycholic acid | 498.475 | 80.042 | 55 | 57.72 | 138 | 0 | 3.75 | 9.5 | 7.68 | 0.15 |
| Taurodeoxycholic acid | 498.475 | 80.042 | 55 | 57.72 | 138 | 0 | 3.75 | 9.5 | 8.1 | 0.2 |
| Cholic acid | 407.25 | 407.25 | 0 | 57.662 | 95 | 77.6 | 5.5 | 10.5 | 9.05 | 0.15 |
| Cholic acid-d4 (internal standard) | 411.388 | 411.388 | 0 | 57.662 | 141 | 100 | 7.5 | 10.5 | 9.04 | 0.15 |
| Ursodeoxycholic acid | 391.338 | 391.338 | 0 | 57.662 | 140 | 100 | 8 | 12.4 | 9.21 | 0.15 |
| Glycochenodeoxycholic acid | 448.25 | 74.089 | 34.4 | 57.662 | 90 | 65.3 | 8.2 | 11 | 9.4 | 0.15 |
| Glycochenodeoxycholic acid-d4 (internal standard) | 452.338 | 73.988 | 38.11 | 57.662 | 98 | 71.4 | 8 | 11 | 9.38 | 0.15 |
| Glycodeoxycholic acid | 448.25 | 74.089 | 34.4 | 57.662 | 90 | 65.3 | 8.2 | 11 | 9.8 | 0.15 |
| Tauroolithocholic acid | 482.338 | 80.03 | 52.05 | 57.662 | 249 | 100 | 8.5 | 11.5 | 10.12 | 0.15 |
| Chenodeoxycholic acid | 391.338 | 391.338 | 0 | 57.662 | 140 | 100 | 8 | 12.4 | 11.04 | 0.15 |
| Chenodeoxycholic acid-d4 (internal standard) | 395.338 | 395.338 | 0 | 57.662 | 126 | 89.8 | 9.5 | 12.2 | 11.03 | 0.15 |
| Deoxycholic acid | 391.338 | 391.338 | 0 | 57.662 | 140 | 100 | 8 | 12.4 | 11.23 | 0.1 |
| Deoxycholic acid-d4 (internal standard) | 395.338 | 395.338 | 0 | 57.662 | 126 | 89.8 | 9.5 | 12.2 | 11.22 | 0.15 |
| Glycolithocholic acid | 432.4 | 74.071 | 32.63 | 57.662 | 112 | 2 | 10 | 12.35 | 11.39 | 0.15 |
| Lithocholic acid | 375.162 | 375.162 | 0 | 83.426 | 119 | 100 | 11.5 | 13.5 | 12.55 | 0.1 |
| Lithocholic acid-d5 (internal standard) | 380.362 | 380.362 | 0 | 83.426 | 119 | 100 | 11.5 | 13.5 | 12.55 | 0.1 |

**Table S3:**
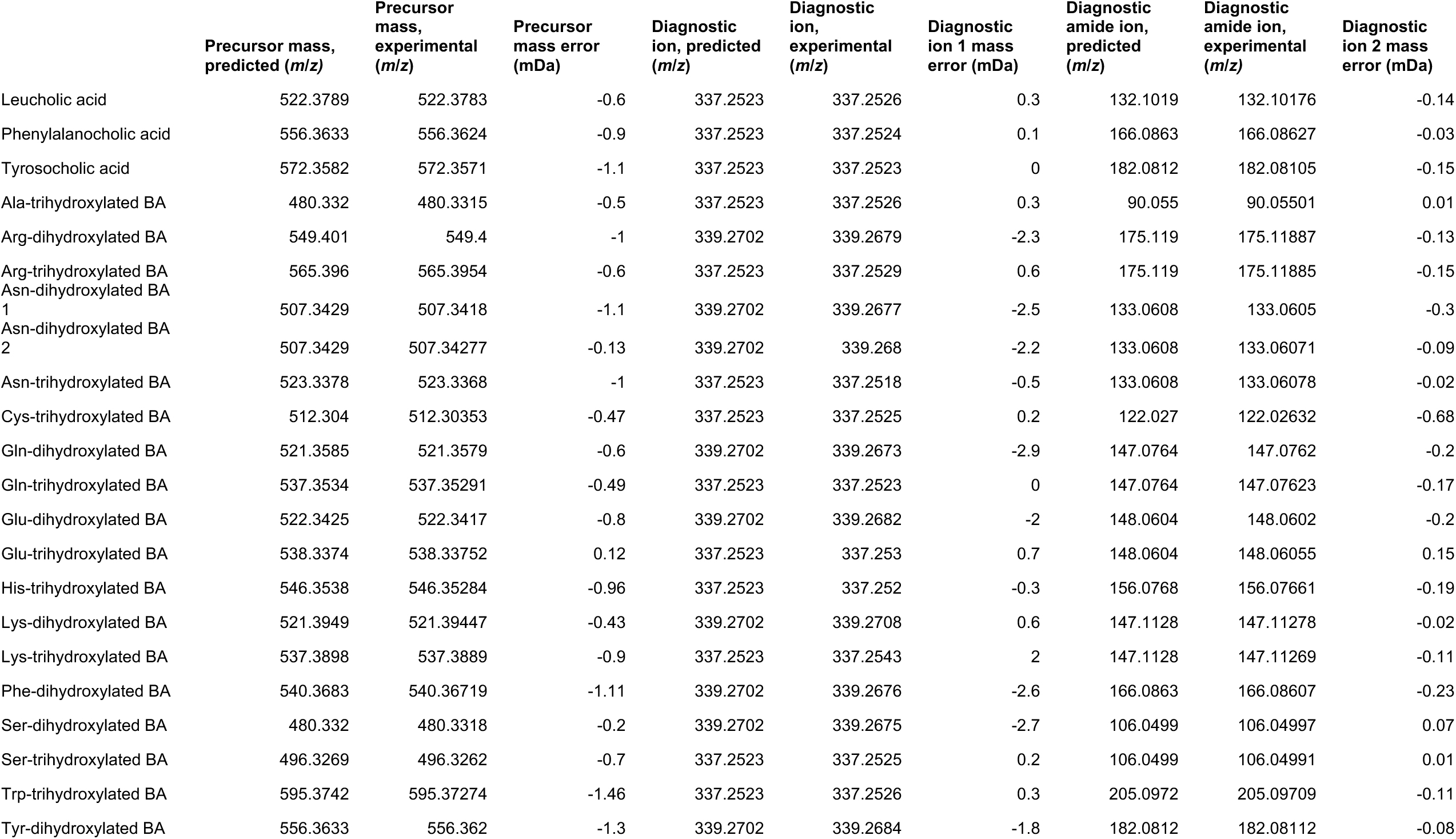
Bile acid conjugate identification from non-targeted analysis. Theoretical and experimental *m*/*z* for precursor mass, and diagnostic fragment ions (MS/MS). Mass error calculated as the difference between the experimental and theoretical *m*/*z*.

## Acknowledgements

The authors thank Laura Symul and Elizabeth Costello for helpful discussions on statistical analyses. This material is based upon work supported by the National Science Foundation under Grant No. 1936687 (to D.S.). The authors acknowledge support from National Institutes of Health Award RM1 GM135102 (to K.C.H.), and the Thomas C. and Joan M. Merigan Endowment at Stanford University (to D.A.R.). J.A.G. is supported by a Stanford University School of Medicine Dean’s fellowship and R.N.C. is supported by an NDSEG graduate fellowship. D.A.R. and K.C.H. are Chan Zuckerberg Biohub Investigators. We acknowledge the Stanford Research Computing Center for computational resources at the Sherlock high-performance cluster. Dedicated to the memory of Yehuda Shalon, who studied bile acids 50 years ago.

