## Supplemental Figures for "Profiling of the human intestinal microbiome and bile acids under physiologic conditions using an ingestible sampling device"

95616, USA
[14]Infectious Diseases Section, Veterans Affairs Palo Alto Health Care System, Palo Alto, CA 94304, USA

*: These authors contributed equally

### Supplemental Figures

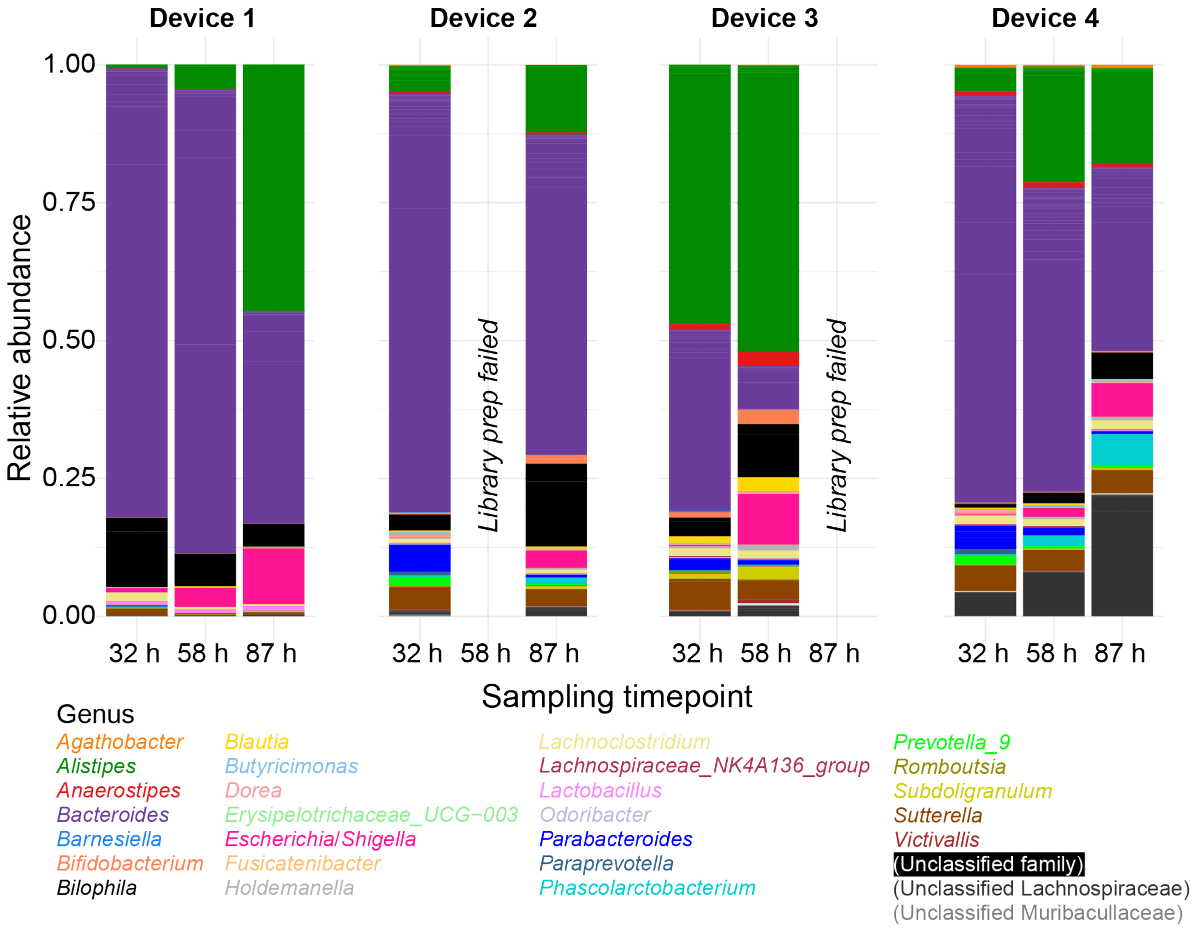


**Figure S1: Changes in intestinal microbiota composition during sample incubation.**

Devices were collected from a bowel movement of a single subject 32 h after ingestion and placed immediately into an anaerobic chamber at 37 °C. Samples were collected from each device immediately (32 h) and again at 58 h and 87 h, and prepared for 16S rRNA gene sequencing. Genus-level relative abundance is shown for all ASVs with read count ≥15 in any single sample.

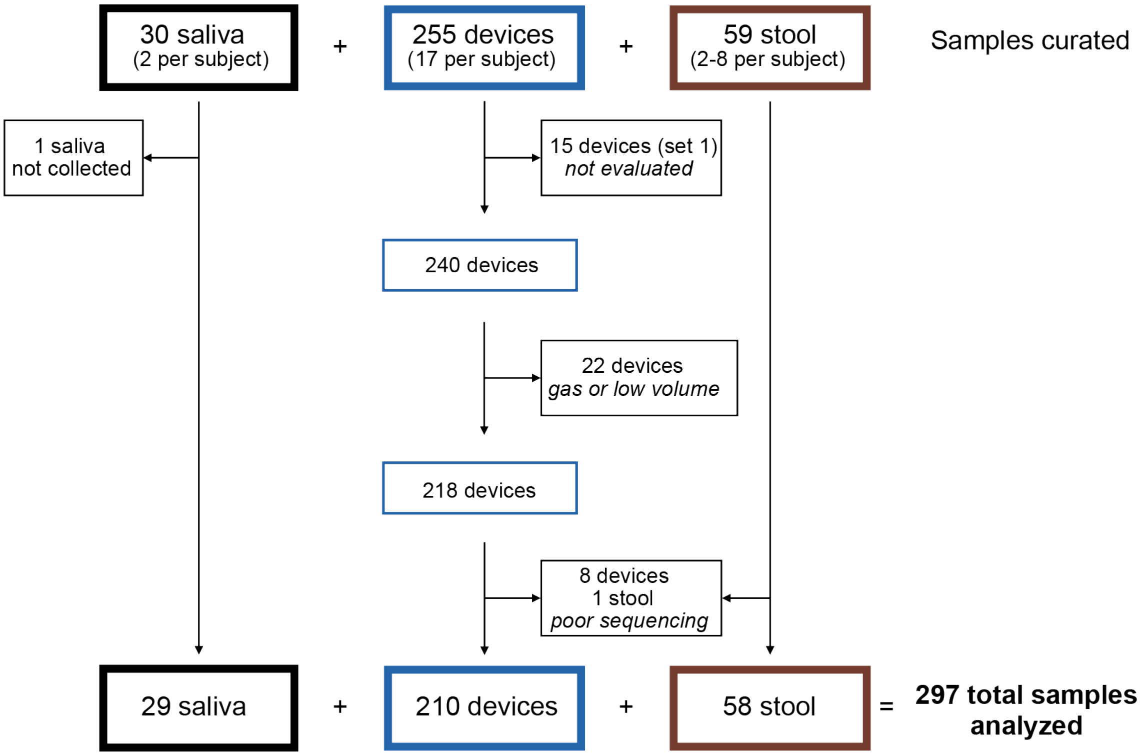


**Figure S2: Sample flow diagram from collection through analysis.**

Subjects (*n*=15) enrolled in the study ingested a total of 17 intestinal sampling devices (set 1 consisted of a single device used as a safety test to ensure safe passage through the intestines). Subjects were also asked to provide 2 saliva samples and collect stool until all intestinal sampling devices were retrieved. Between 2-8 stool samples from each subject were used for analysis.

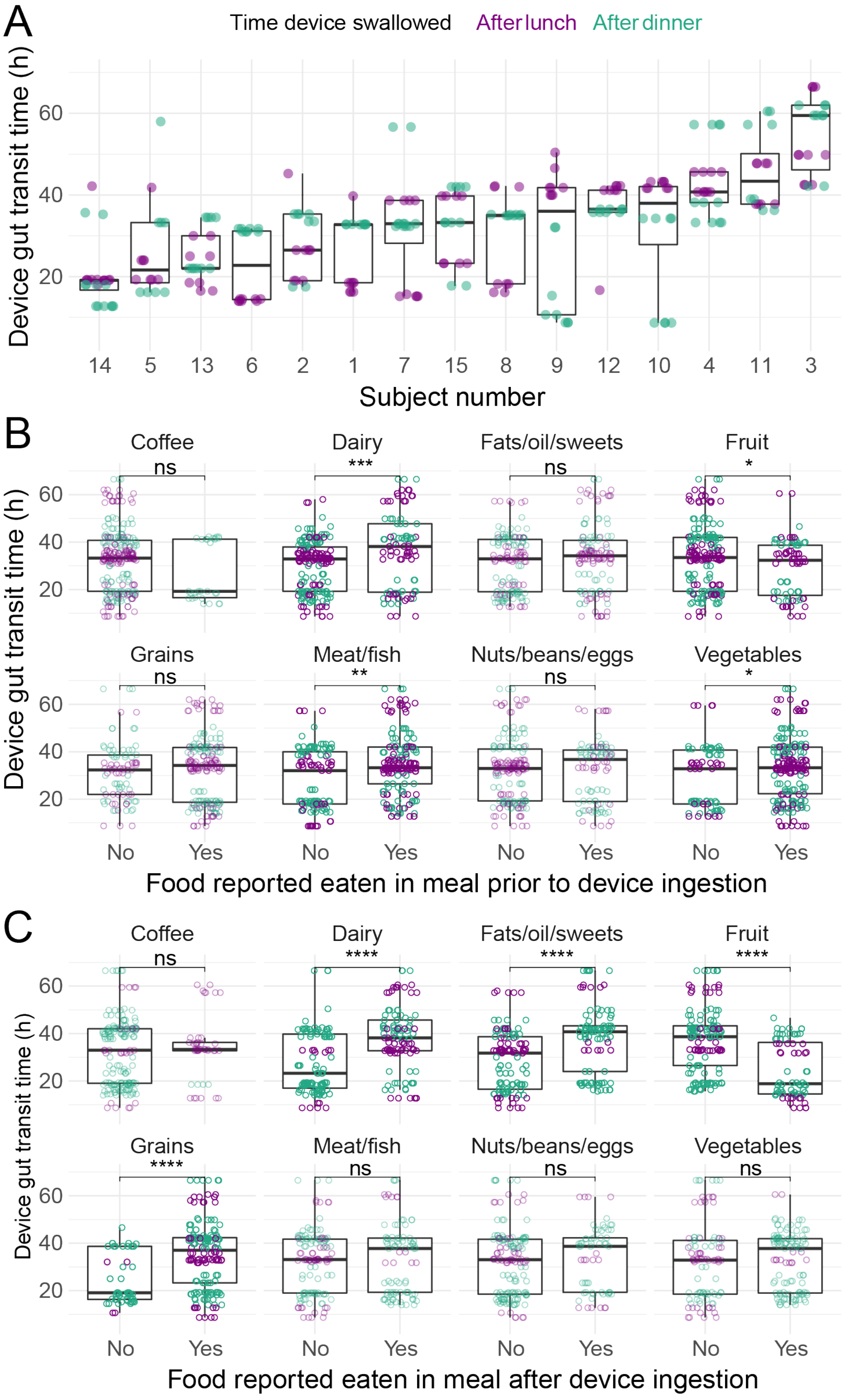


**Figure S3: Total gut transit time of devices varied across subjects and diets.**

A) Device gut transit time was variable across subjects. Some subjects displayed differences in transit time dependent on the time of day the device was ingested.
B) Device gut transit time varied according to certain types of food consumed in the meal prior to device ingestion (i.e., the food with which the devices presumably transited into the small intestines).
C) Device gut transit time varied according to the type of food consumed in the meal after capsules were swallowed (i.e., the food that likely influenced gut motility while devices were passing through the large intestines).

Boxplots show the median, 25$^{th}$, and 75$^{th}$ quartiles. ns: not significant, *: $p \leq 0.05$, **: $p \leq 0.01$, ***: $p \leq 0.001$, ****: $p \leq 0.0001$, Bonferroni-corrected Wilcoxon rank sum test.

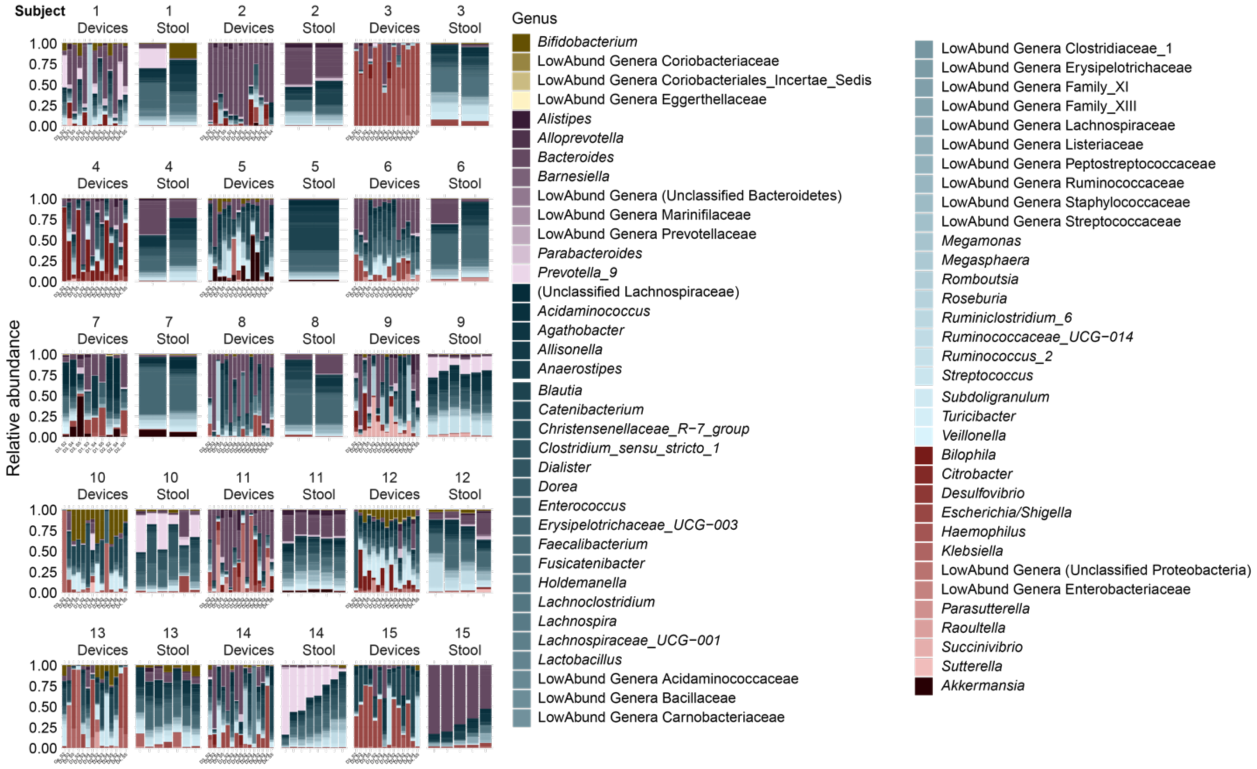


**Figure S4: A wide variety of microbial taxa were detected across subjects in intestinal and stool samples.**

Relative abundance of ASV reads mapped at the genus level are shown for each sample by subject. ASVs that did not have a read count of ≥3 in 5% of samples were ignored. Genera that were not detected at ≥5% in one sample were lumped into family categories for clarity (LowAbund Genera = low abundance genera of a family). Taxa are colored by phylum (Actinobacteria, yellow; Bacteroidetes, purple; Firmicutes, blue; Proteobacteria, red; Verrucomicrobia, black). D# = device type; S# = set.

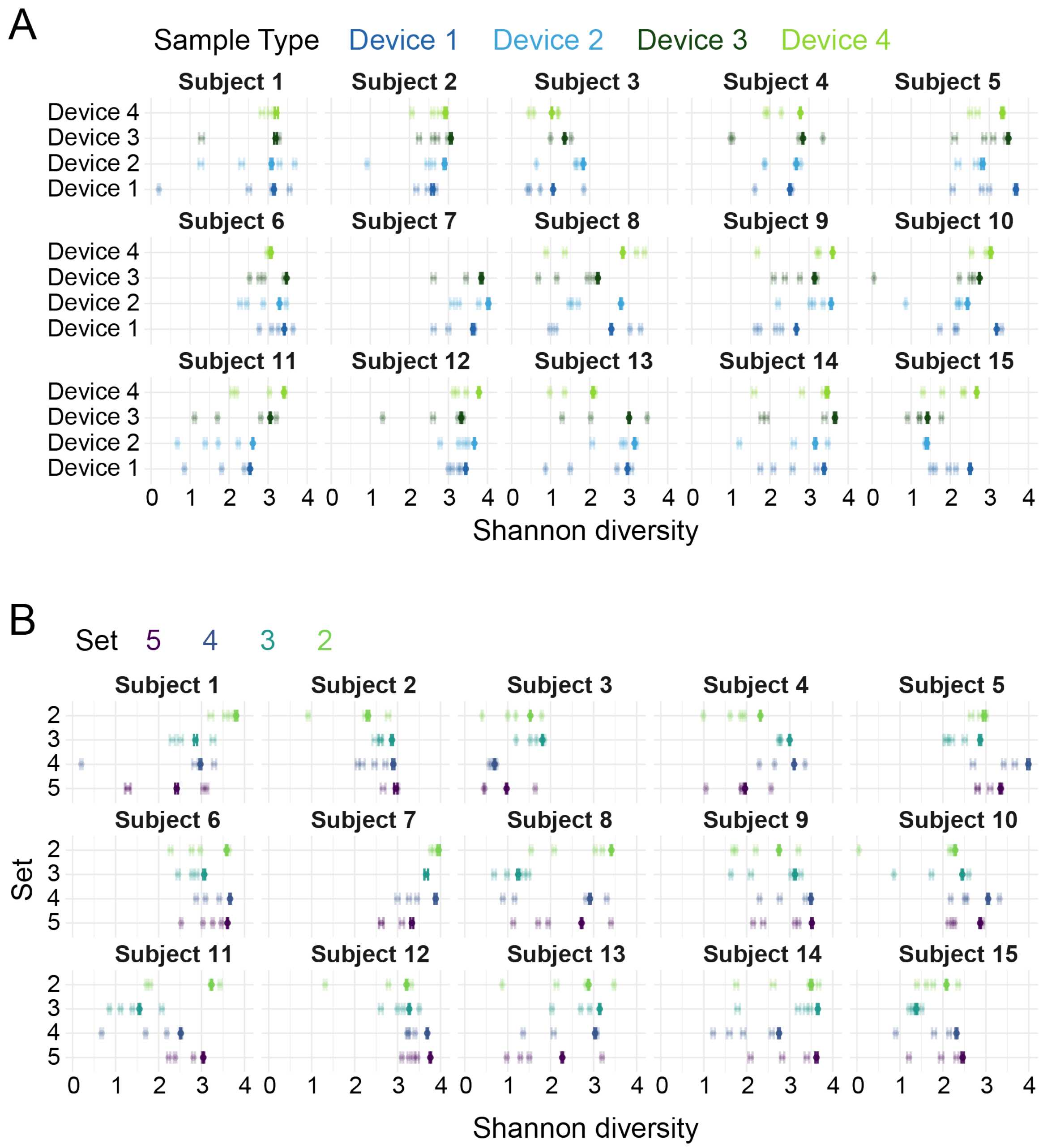


**Figure S5: Alpha and gamma diversities implicate a temporally and spatially heterogeneous intestinal tract.**

A) The Shannon diversity of individual devices (alpha diversity, lighter points) was generally lower than that of all devices of a certain type (gamma diversity, bold points), indicating high temporal variation.

B) Greater overall diversity in a set of devices swallowed at the same time show spatial patchiness. The Shannon diversity of individual devices (alpha diversity, lighter points) was generally lower than that of all devices from a single set (gamma diversity, bold points), indicating high spatial variation.

Each subject (#1-#15) is shown separately. To ensure equal read depths for accurate comparisons, all intestinal samples were rarefied to the minimum sequencing depth of any device from that subject. Mean values and 95% confidence intervals for alpha and gamma diversity estimates were obtained by repeating the rarefaction procedure 1000 times.

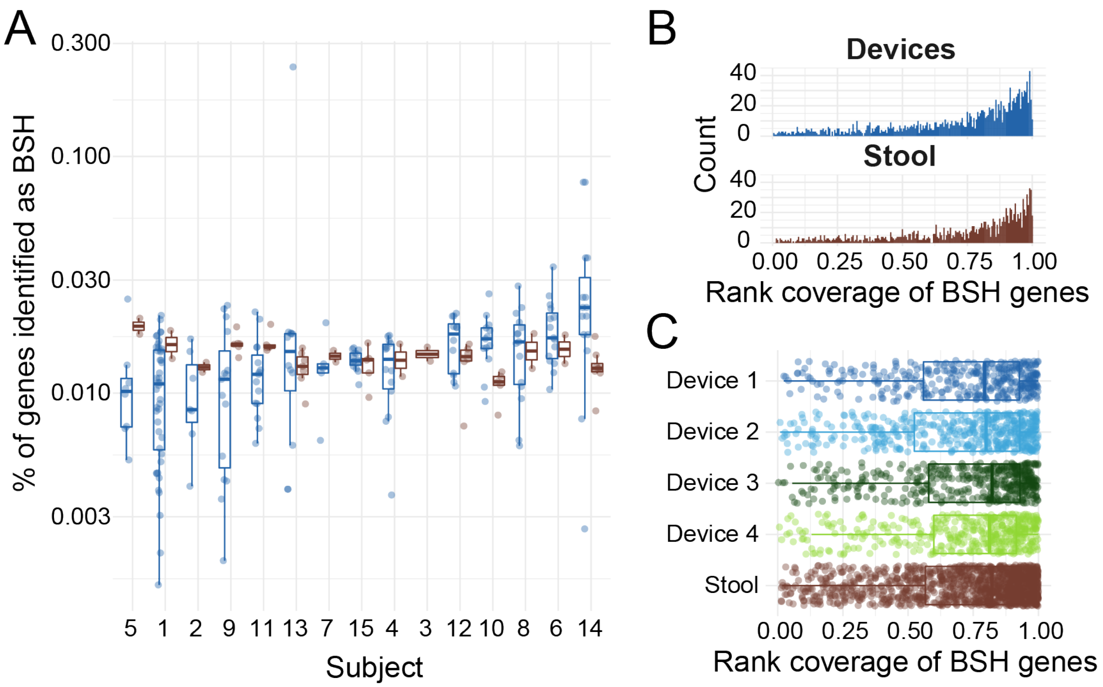


**Figure S6: Microbial bile salt hydrolase genes exhibited similar abundance and diversity in intestinal and stool samples.**

A) Open reading frames identified as bile salt (cholylglycine) hydrolase (BSH) enzymes via a hidden Markov model (HMM) search, normalized by the total number of open reading frames detected in the sample.

B) The distribution of rank coverage of *bsh* genes was similar between intestinal and stool samples.

C) Rank coverage of *bsh* genes in devices of each type and in stool are similar.

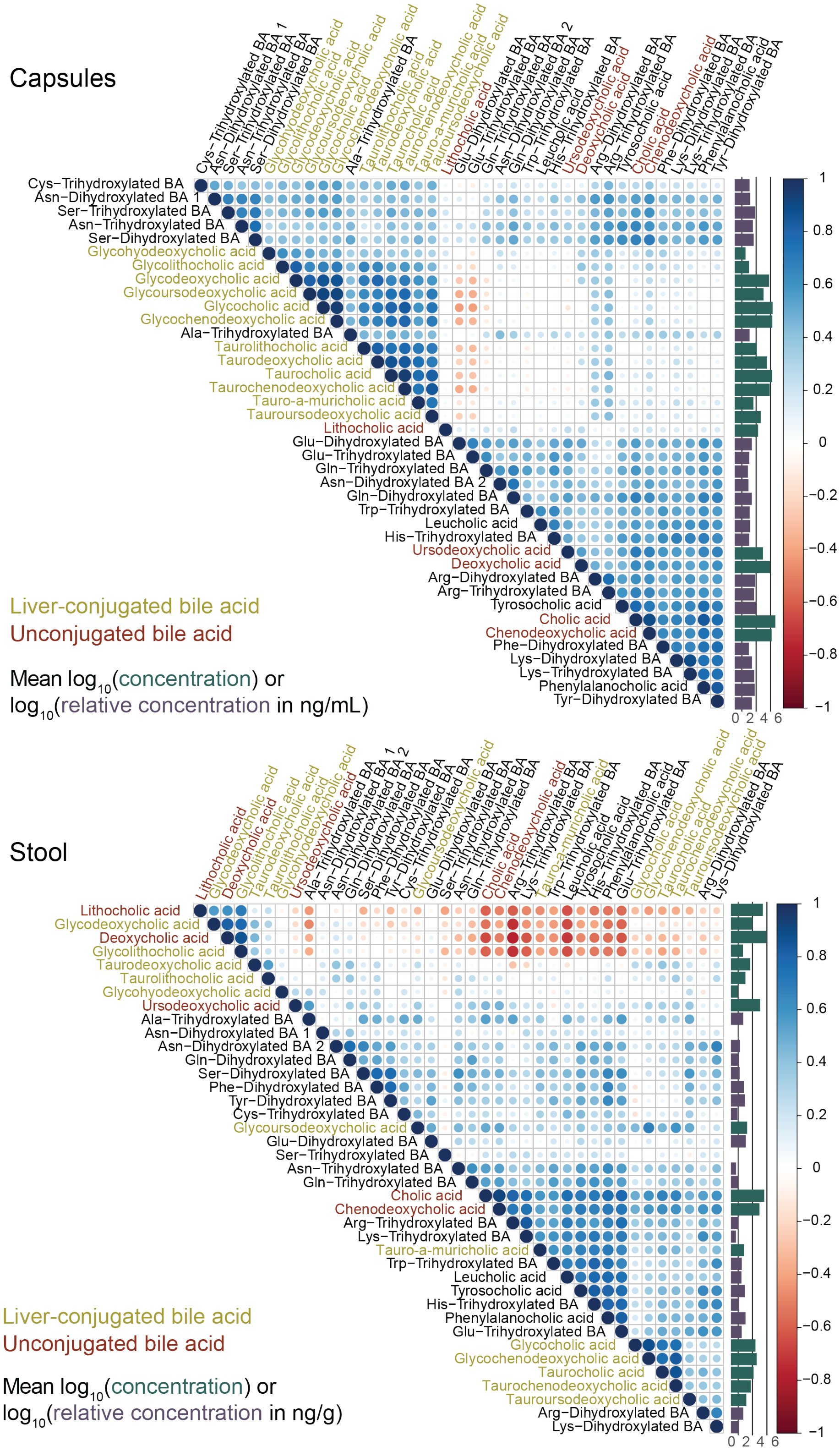


**Figure S7: Devices capture distinct bile acid profiles along the intestinal tract compared with stool.** Correlations between $\log_{10}$(concentration) of every bile acid chemical structure (correlations for coarser groupings are shown in Fig. 3E) in intestinal (top) and stool (bottom) samples. Average $\log_{10}$(concentration) of a bile acid in each sample type is shown in the bar plot on the right.

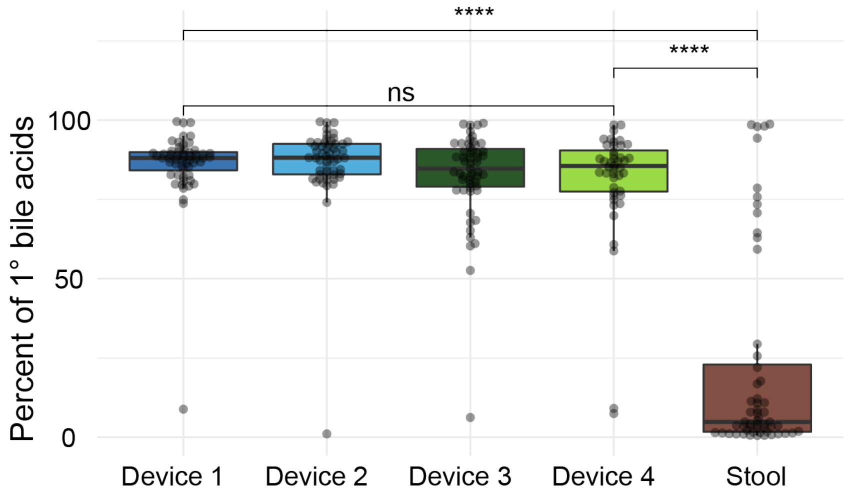


**Figure S8: Percentage of primary (hydroxylated) bile acids was similar across device types and was lower in stool compared with intestinal samples.** Boxplots show the median, 25$^{th}$, and 75$^{th}$ quartiles. ns: not significant, ****: $p \leq 0.0001$, Bonferroni-corrected Wilcoxon rank sum test.

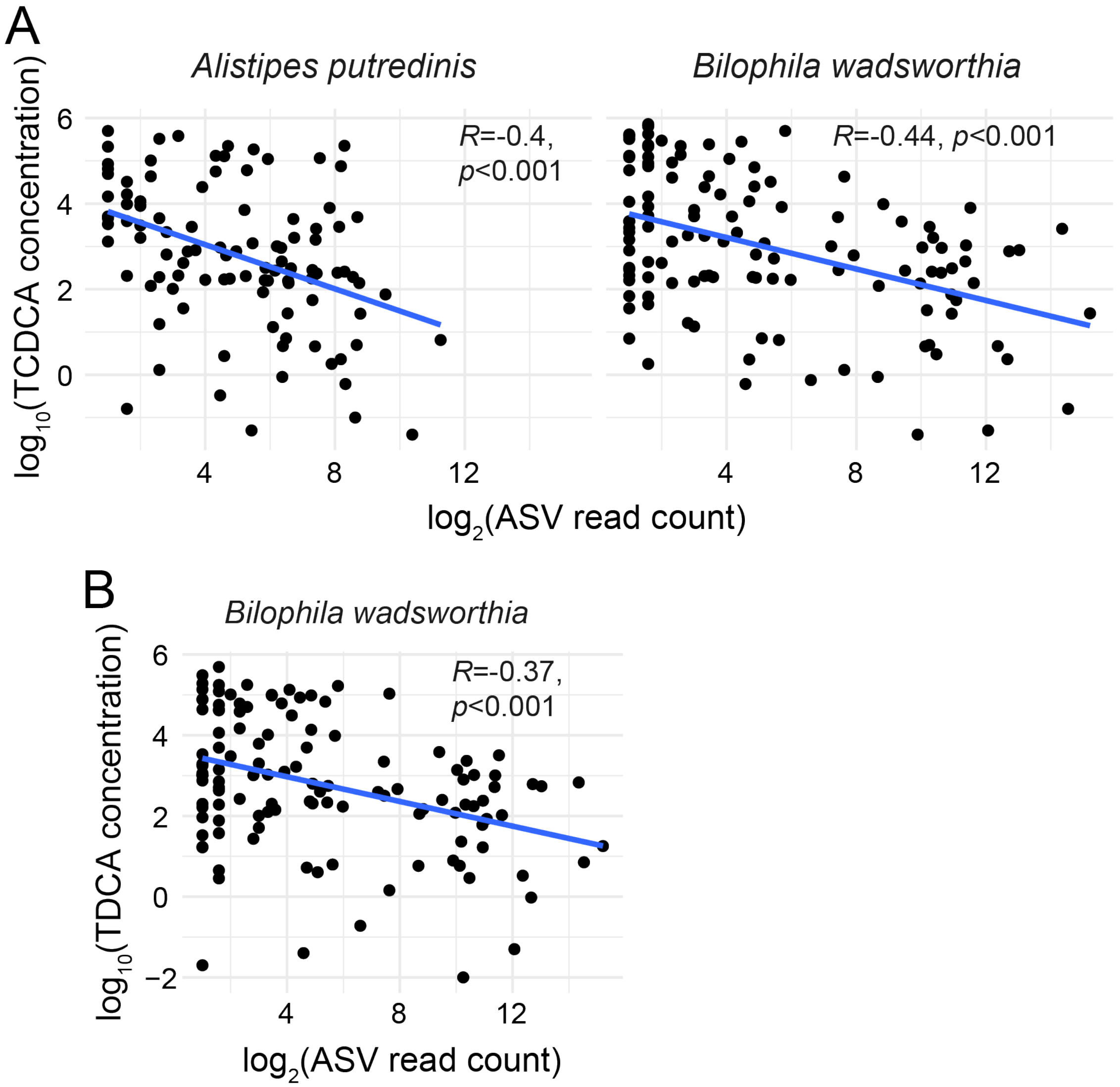


**Figure S9: Taxa negatively correlated with TCA concentration were also negatively correlated with the concentration of other taurine-conjugated bile acids.**

A) The $\log_2$(ASV count) of *Alistipes putredinis* and *Bilophila wadsworthia* was negatively correlated (Pearson, Benjamini-Hochberg-corrected) with the $\log_{10}$(concentration) of primary bile acid taurochenodeoxycholic acid (TCDCA) across all intestinal samples.

B) The $\log_2$(ASV count) of *B. wadsworthia* was negatively correlated (Pearson; Benjamini-Hochberg-corrected) with the $\log_{10}$(concentration) of taurodeoxycholic acid (TDCA), the secondary bile acid formed by dehydroxylation of TCA, across all intestinal samples.

Only ASVs with $p<0.01$ after a Benjamini-Hochberg correction are shown. Significant negative correlations between the $\log_2$(ASV count) of any taxon and glycine-conjugated bile acids aside from GCA were not observed.

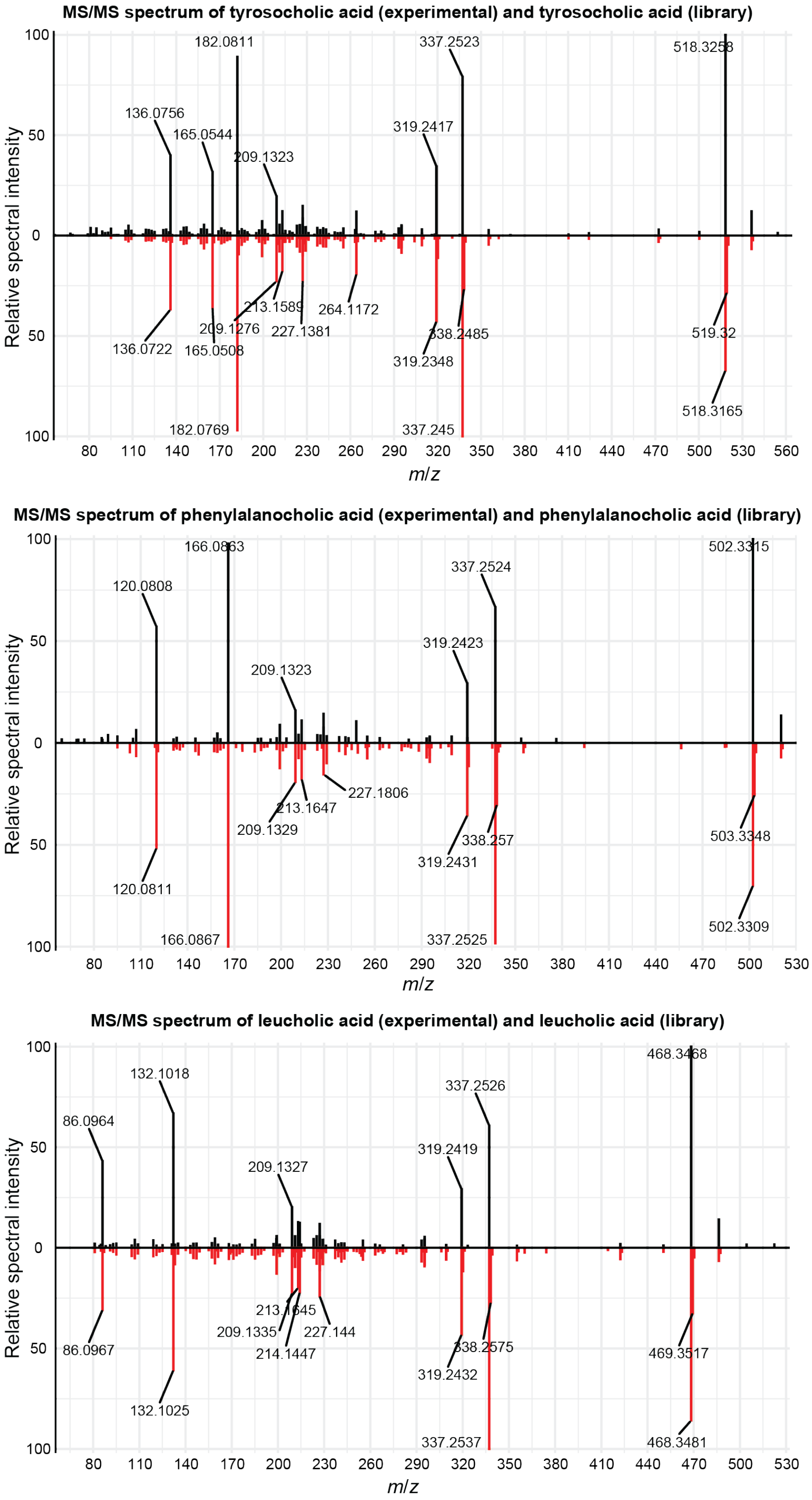


**Figure S10: Microbially conjugated bile acid identification.** Head-to-tail matches of experimental (top) to library (bottom) spectra from bile acids conjugated to tyrosine, phenylalanine, and leucine.
